# Distributed learning across fast and slow neural systems supports efficient motor adaptation

**DOI:** 10.1101/2025.06.01.657238

**Authors:** Leonardo Agueci, N Alex Cayco-Gajic

## Abstract

Adaptation is a fundamental aspect of motor learning. Intelligent systems must adapt to perturbations in the environment while simultaneously maintaining stable memories. Classic work has argued that this trade-off could be resolved by complementary learning systems operating at different speeds; yet the mechanisms enabling coordination between slow and fast systems remain unknown. Here, we propose a multi-region distributed learning model in which learning is shared between two populations of neurons with distinct roles and structures: a recurrent ‘controller’ network which stores a slowly evolving memory, and a feedforward ‘adapter’ network that rapidly learns to respond to perturbations in the environment. In our model, supervised learning in the adapter produces a predictive error signal that simultaneously tutors consolidation in the controller through a local plasticity rule. Our model offers insight into the mechanisms that may support distributed computations in the motor cortex and cerebellum during motor adaptation.

## Introduction

Flexible, goal-directed behavior requires a balance between adaptation and stability of motor representations. Organisms interact with an ever-changing environment, which can transform how neural activity is mapped onto behaviour [1–3]. To reflect these changes, the brain must store and update memories representing the task [4]. If a given perturbation represents a permanent change, the memory should be adapted as a result. However, whether a perturbation will be persistent or transient is not known a priori. Adapting too early to a temporary change would needlessly overwrite the previous motor memory, raising the question: how do organisms balance a trade-off between stability and adaptability of motor memories?

A resolution to this dilemma posits the existence of dual learning processes, one slow and one fast, which jointly drive motor behavior [5]. According to this hypothesis, a fast system adapts rapidly to external changes in the environment, while a slow learner maintains a stable memory that is progressively updated. An interplay between slow and fast learning processes is at the heart of key theories of human behavior in sensorimotor adaptation [5, 6], habit formation [7, 8], and episodic memory [9, 10]. However, despite behavioral evidence for simultaneous, multiple-timescale learning [5, 6, 11, 12], the identities of the slow and fast learners, as well as the computational mechanisms enabling their coordination, remain a subject of debate.

Motor learning is known to rely on the synergistic efforts of multiple brain areas, each characterized with its own specific function, circuitry, and learning rules [13]. Among those structures, the motor cortex and cerebellum stand out as prime candidates for the slow and fast learners [14]. Indeed, the motor cortex has been shown to store stable motor memories in its recurrent dynamics [15–18]. Conversely, the cerebellum is known to be involved in adaptation to environmental changes through rapid, error-based plasticity [19–23]. While traditionally studied in isolation, recent work has highlighted the interactions between these regions in goal-directed behaviors [24], including motor planning [25] and motor learning [26, 27], where the cerebellum may provide strong input drive that can shape motor cortical activity [28–30]. Recent modeling has proposed a replay-like mechanism that allows consolidation of the cerebellar contribution to the cortex [31]. However, the mechanisms by which the cerebellum and motor cortex could cooperate as *simultaneous* learning systems remain unknown.

To address this gap, we introduce a multi-region neural network model that can implement a two-timescale distributed learning strategy. Our model consists of an intercoupled recurrent *controller* network and a feedforward *adapter* network, which together form an internal model that jointly drives motor behavior. The controller stores a stable memory of the baseline task in its recurrent dynamics, while the adapter rapidly learns to respond to perturbations in the environment using supervised learning based on external sensorimotor errors. The multi-region circuitry is constructed so that the adapter learns to predict the internal cortical error which can be used in three ways: to correct cortical dynamics following principles of feedback control, to directly correct cortical output itself, and to tutor slow consolidation of the new memory using a local plasticity rule. Remarkably, all three of these goals can be achieved simultaneously by using the same signal from the adapter. Learning occurs simultaneously in both regions, without the need for a separate consolidation phase or replay.

The resulting model builds upon classic theories of motor adaptation and learning [13, 32–36], by adding a supplementary role for the cerebellum as both a tutor and driver of cortical activity. By leveraging established theories of multiple learning timescales in the brain [5, 9, 11], our hypothesis sheds light on recent evidence of the cerebellum’s influence on cortical activity [28–30] and the involvement of the cerebello-spinal projection during motor learning [37]. Moreover, it provides strong predictions for inactivation experiments that could clarify the cerebellum’s role in motor learning and adaptation.

## Results

### The distributed learning hypothesis

We consider a recurrent neural network (the *controller*, representing the motor cortex) which executes a motor plan encoded in its upstream inputs, such as a goal-directed reach [38]. The controller’s recurrent weights store a memory of the dynamics needed to perform the movement. However, downstream changes in the actuators or environment can change the mapping between neural activity in the controller and the hand trajectory (e.g., in the visuomotor or force field adaptation tasks [2, 3, 27]). Such changes effectively perturb the target trajectory that must be performed by the controller (Figure 1a, see Methods). We assume that the upstream input to the controller remains fixed, so that it must use the sensorimotor error accumulated over its enacted hand trajectories to guide trial-by-trial learning. How does the controller reorganize its internal dynamics over trials to compensate for the perturbation?

**Figure 1:**
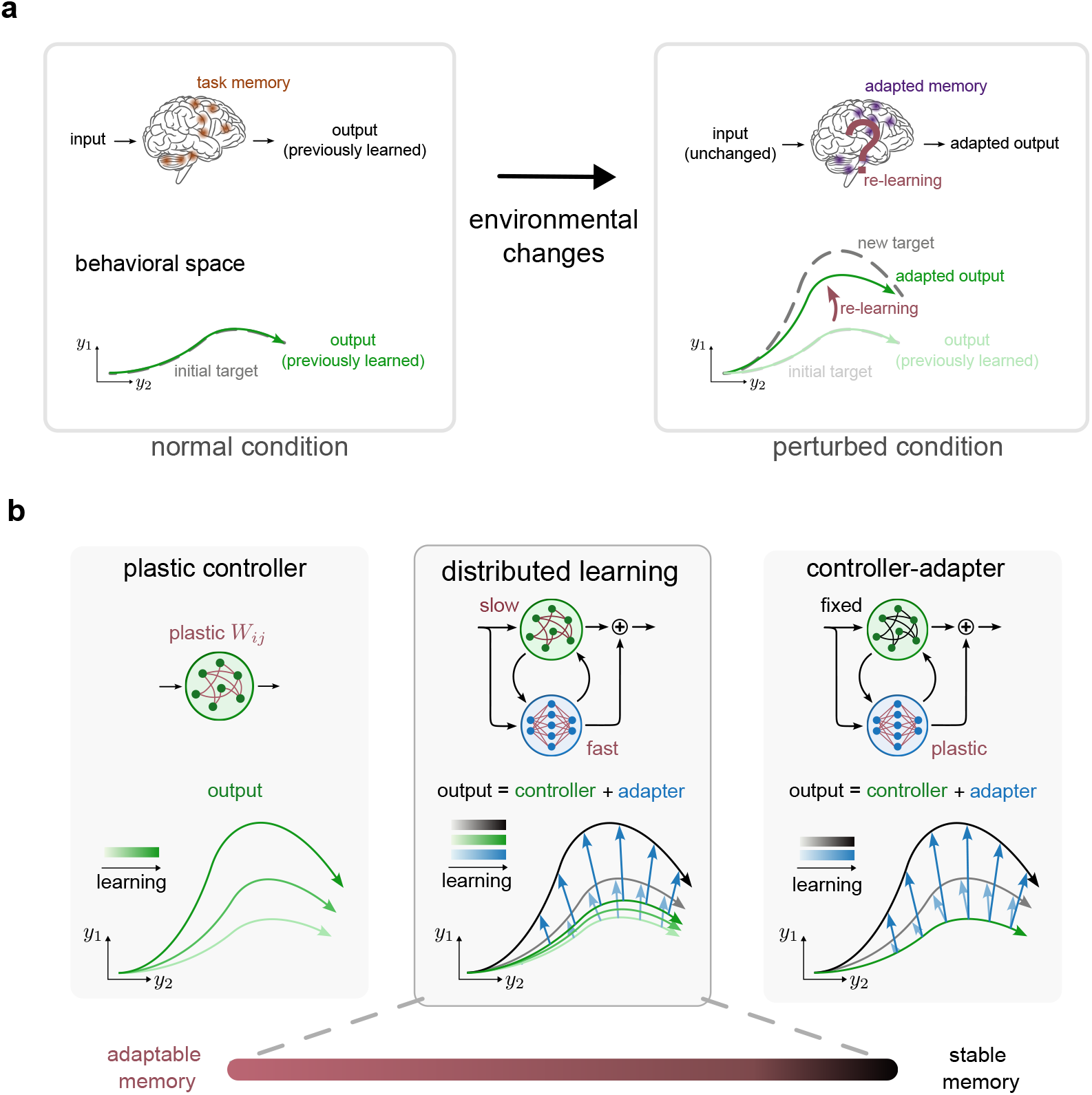
Three hypotheses for how recurrent dynamics adapt previously acquired skills. **a,** Neural populations perform tasks by generating a specific output in response to an upstream input signal. Perturbations in the environment modify the effective target output without changing upstream input. The network must adapt its internal dynamics so that the output moves from the initial target to the new target. **b,** Different hypotheses for adapting to perturbations. Left: A recurrent neural network (RNN) with plastic weights (indicated in red). The RNN output gradually converges to the new target. Right: An RNN (the *controller*) with non-plastic weights stores a stable memory of the original output. Instead, a secondary population (the *adapter*) learns the residual between the controller’s output (corresponding to the initial target) and the new target. Center: We propose a two-timescale distributed learning framework in which both the controller and the adapter networks are plastic: the adapter rapidly learns to correct the controller’s output while simultaneously tutoring the controller to consolidate the updated memory into its dynamics. Over a long timescale, the controller output converges towards the new target. Depending on the relative learning rates in the controller and adapter, the resulting model may be positioned anywhere along the adaptability-stability spectrum.

A simple hypothesis is that the RNN simply rewires its recurrent connections to account for the perturbation [39]. In this *plastic controller* hypothesis, the output is gradually modified over trials until it matches the new target (e.g., through gradient descent; Figure 1b, left). However, this could pose a significant problem in the face of transient perturbations, as the controller would need to relearn the original dynamics for the baseline task once the perturbation has been abolished [40]. A second hypothesis, inspired by classic work on adaptive feedback control [41], proposes that the controller instead maintains a stable memory of the original task, while relegating any modification to a secondary population, the *adapter* network. In this scenario, the adapter learns the residual necessary to correct the output of the controller (Figure 1b, right). These two hypotheses exemplify two views on learning at opposing ends of a stability-adaptability trade-off: the controller is either entirely plastic or entirely stable.

Building on this perspective, we propose a third hypothesis in which the controller and the adapter are both plastic and work together as a single *distributed learning* system: the adapter rapidly minimizes the behavioral error, while the controller slowly consolidates the new memory into its internal dynamics (Figure 1b, center). This allows the system to implement a two-timescale learning strategy in which the memory encoded in the controller remains stable on a short timescale, but can be updated on a long timescale in response to persistent changes [5]. The short-term stability of the controller memory provides a baseline upon which the adapter can learn a prediction of the controller’s internal error signal. Conversely, the adapter is able to use this predictive error to tutor the controller to consolidate the memory of the perturbed task.

In the remainder of the manuscript, we will systematically demonstrate how such a distributed learning system can be constructed in four steps. First, we pretrain a recurrent controller network that uses structured error feedback as an input drive, allowing it to correct its dynamics without modifying its recurrent weights [42, 43]. This ensures that the controller is able to partially generalize to external perturbations without requiring additional plasticity that would overwrite the memory of the baseline task. Second, we replace the actual error feedback with a predicted error learned by the adapter, forming a multi-region network in which only the adapter is plastic. Here, the adapter module simultaneously drives the controller’s dynamics and fine-tunes its output. Third, we introduce slow, gradient-based plasticity in the controller, that allows it to consolidate the memory of the perturbed task, forming a two-timescale distributed learning system. This is achieved by optimizing the recurrent weights based on the decoded hand trajectory of the controller alone, without any contribution from the adapter. This third step therefore validates the architectural choices necessary to ensure coordinated learning between the two systems. In the final step, we switch to a local plasticity rule in the controller [42, 43] which is mediated by the adapter’s feedback. With this local learning rule, the adapter feedback directly tutors consolidation of the updated memory in the controller.

### Pretraining a feedback controller

We start by building a controller network that can perform feedback control by integrating an externally provided error signal. To this end, we pretrained RNNs with backpropagation to perform a center-out reaching task (Figure 2a,b). Taking inspiration from motor control theory [4, 36, 41], we assumed that the RNN could only control the behavioral output **y**(*t*) indirectly, by producing a motor command ***θ***(*t*) which is fed into a nonlinearity *P* (called the *plant*; Figure 2b). The goal of the controller is not to generate the target output **y**\*(*t*), but rather to produce the target motor command ***θ***\*(*t*) = *P* ^−1^(**y**\*(*t*)) when provided with a reference trajectory (Methods). As such, the controller is a neural network implementation of an inverse model [36].

**Figure 2:**
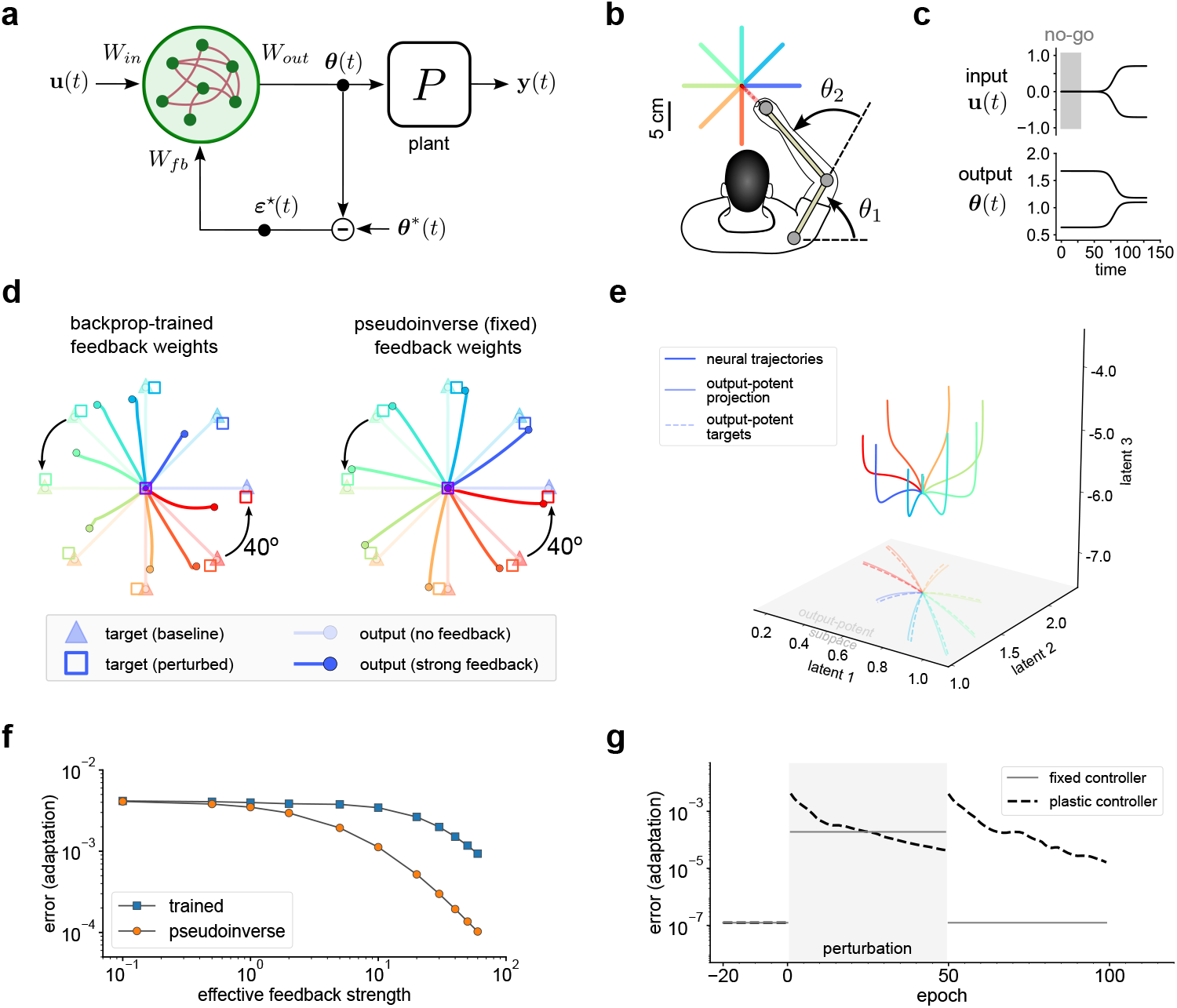
Online correction of RNN dynamics requires structured error feedback for generalization to perturbations. **a,** A feedback RNN that generates the motor command signal ***θ***(*t*) necessary to control a downstream motor plant *P*. The controller receives feedback in the form of the motor command error signal: ***ε***⋆(*t*) = ***θ***\*(*t*) − ***θ***(*t*). **b,** Schematic of the center-out reaching task and the motor plant. The plant is modeled as the nonlinear mapping from the shoulder and elbow angles to the 2-dimensional hand position (colored trajectories). **c,** Example input and motor command for one center-out trajectory. The input **u**(*t*) consists of a binary no-go signal (grey) and the 2-dimensional reference trajectory of the movement in Euclidean coordinates (top). The RNN generates the target motor command that will drive the plant to reproduce the target trajectory. **d,** Performance of trained RNNs in a perturbation task in which the targets are rotated by 40 degrees while inputs **u**(*t*) and all weights remain fixed. RNN feedback weights were either trained with backpropagation (left), and or fixed to the pseudoinverse of the readout weights: 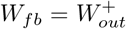 (right). Pseudoinverse feedback weights led to better generalization to large perturbations. Trajectories are shown for both no-feedback and strong-feedback cases (*κ* = 0 vs. *κ* = 30, see Methods). **e,** Strong error feedback corrects dynamics by realigning trajectories to the target in the output-potent subspace. Example neural trajectories visualized in a three-dimensional subspace (corresponding to the output-potent dimensions 1 and 2, and the first principal component). **f,** Error in the perturbation task as a function of the effective feedback strength (see Methods) for trained (blue) and fixed pseudoinverse (orange) feedback weights. **g,** Performance of the fixed feedback controller (grey line) compared to a standard RNN with trained weights (plastic controller, dashed black line) during a 40 degree perturbation and washout. The fixed controller is able to only partially correct its dynamics during the perturbation, but can immediately revert to the original task.

Based on this premise, the controller received two sources of input: upstream input **u**(*t*) (consisting of a go/no-go cue and the reference trajectory, presumably from higher-order cortical areas [38]; Figure 2c), and a motor command error signal ***ε***⋆(*t*) = ***θ***\*(*t*) − ***θ***(*t*). For now, we will assume that the error signal is provided; we will address where this signal might originate in the following section. During pretraining, the principal dynamic modes of the recurrent weights showed an increased alignment with the feedback weights (Supplementary Figure 1d,e).

After pretraining the controller on the baseline task, we next tested whether it could generalize its performance to an external perturbation without any additional plasticity [43]. We rotated the targets by a fixed angle, resulting in new target hand kinematics 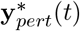 and corresponding new target motor commands 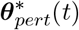 (equivalent to a visuomotor rotation; see Methods). This resulted in a change in the error observed by the controller, while the rest of the model (all weights and the upstream input **u**(*t*)) remained fixed. Thanks to the error feedback, the controller was able to partially redirect its behavioral trajectories (Figure 2d, left). However, this modest correction was insufficient to attain the new target endpoints without additional training, even for small perturbations (Supplementary Figure 2a,b).

The ability of the controller to generalize can be improved by considering both the amplitude and the structure of the error feedback signal. First, previous work has suggested that strong error feedback can force RNN dynamics to follow a reference trajectory [42, 44]. Indeed, increasing the strength (or gain) of the feedback weights led to partial improvements in generalization without any further optimization (until a critical value after which the system loses stability; Supplementary Figure 3). Second, the controller can make more efficient use of the error signal by structuring its feedback weights to be aligned with the output-potent dimensions of the neural space; this way, the error feedback can directly correct the controller’s population activity along the dimensions that lead to an improvement in performance (Supplementary Figure 4). This can be ensured by fixing the feedback weights to the pseudoinverse of the readout weights: 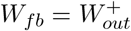.

**Figure 3:**
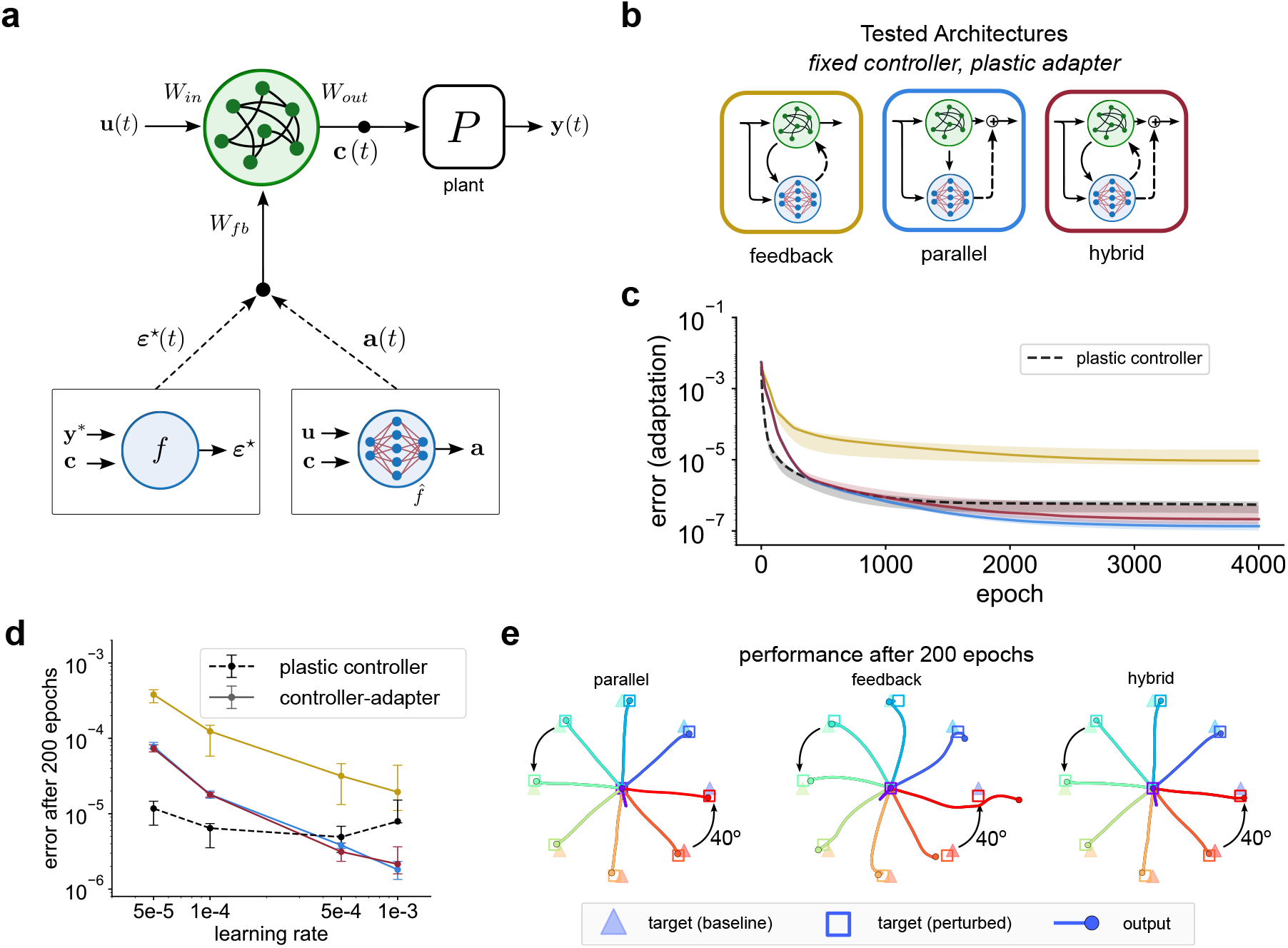
Rapid correction of controller output and dynamics through error approximation. **a,** Schematic depicting the role of the adapter network in predicting the controller’s internal error, ***ε***⋆(*t*). Bottom left: ***ε***⋆(*t*) is a function *f* of the target trajectory **y**\* and the controller output **c**(*t*) (an efference copy from the controller; see Methods). Bottom right: Instead, a feedforward adapter network can be used to approximate ***ε***⋆(*t*) as a function 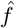 of the internally available signals: **u**(*t*) and **c**(*t*). This prediction is learned over trials: after each reach, the behavioral error is used to refine the plastic adapter weights (red), improving the accuracy of the predicted internal error. **b,** Schematics of the three tested controller-adapter architectures. Dashed lines represent the adapter output pathways that distinguish the three architectures (feedback to the controller, projection to the plant, or both). Solid lines represent the pathways that are present for all architectures: a copy of **c**(*t*) and **u**(*t*), needed for the adapter’s function approximation. In all models, the controller weights are frozen (black) after being pretrained to the baseline task. Adapter weights (red) are trained with backpropagation during the perturbed task. **c,** Mean-squared error (MSE) during adaptation to a 40° rotation of center-out task. The three different controller-adapter architectures (color-coded according to **a**) are compared to a plastic controller. Solid lines, mean; shaded areas, min-max over 5 random seeds. The feedback architecture has the poorest performance. Results shown for adapter learning rate 1e-4. **d,** Average MSE after 200 epochs for different architectures using different learning rates. **e,** Performance of the three controller-adapter architectures after 200 training epochs on the adaptation task. Both parallel and hybrid models show near-total adaptation, compared to the feedback architecture. Trajectories were extracted from the same trainings as in **c**.

**Figure 4:**
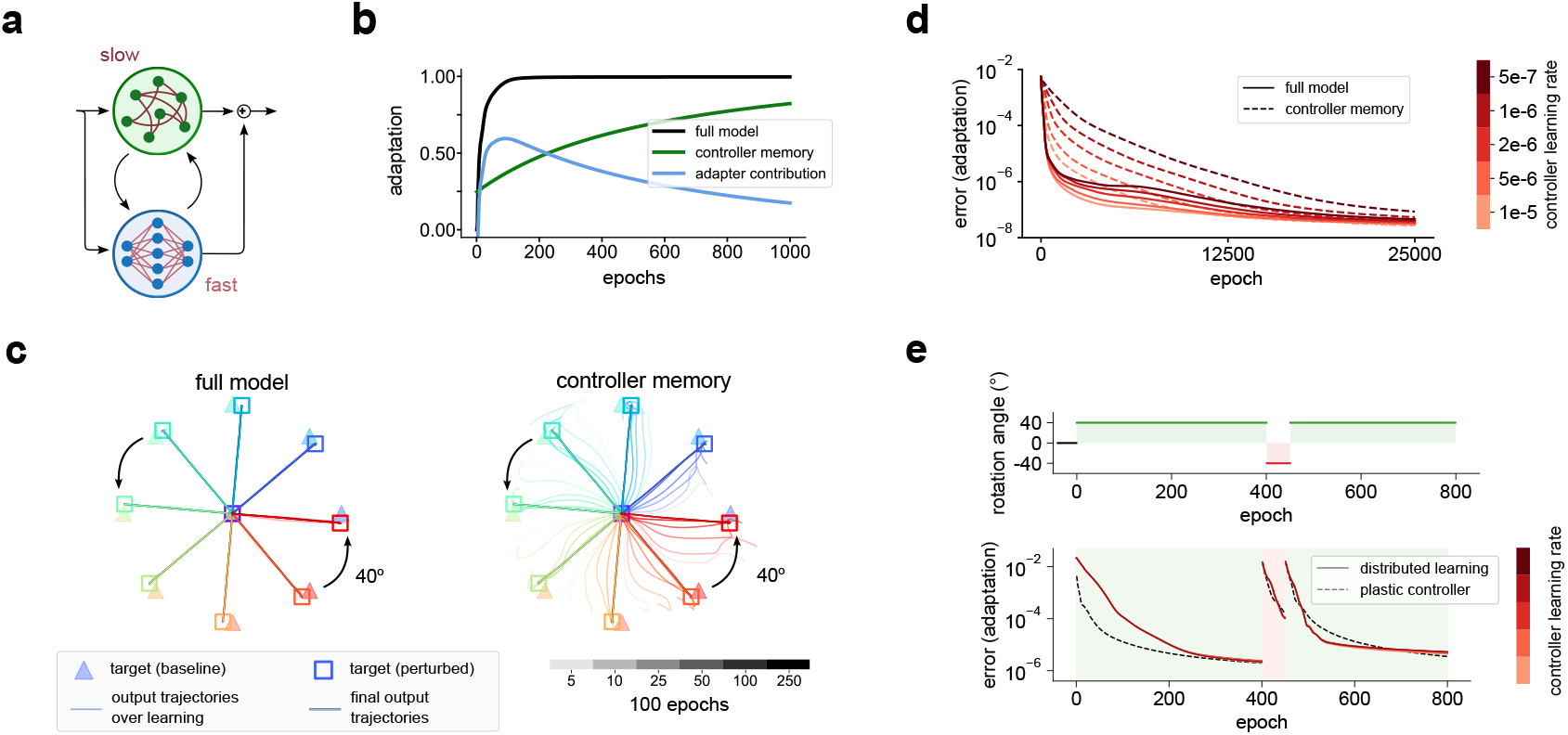
Simultaneous rapid adaptation and slow memory consolidation through distributed, two-timescale learning. **a,** Schematic of the hybrid architecture implementing distributed learning. Both modules were trained with backpropagation, with different learning rates for the controller (green) and the adapter (blue). The adapter is trained to minimize behavioral error, while the controller is trained to minimize the error of it stored memory (Methods). **b,** Adaptation level during the perturbed task, showing the separate contributions of the controller and adapter (Methods). The adapter displays a rapid-onset transient contribution while the memory in the controller is only slowly updated. Controller learning rate: 5e-7. Adapter learning rate: 1e-4. **c,** Evolution of the behavioral trajectories during adaptation to the perturbed task (same simulations as in **b**). Left: The full model almost perfectly learns the task after 500 trials and maintains it for the rest of learning. Right: probe trials in which the behavioral output was decoded from the controller only (i.e. with the adapter silenced), showing that the memory encoded in the controller’s intrinsic dynamics slowly converges to the new targets. Single trajectories are color coded for different target endpoints, and brightness indicates the epoch during learning. **d**, Effects on the speed of adaptation and consolidation during washout for different consolidation speeds (i.e., varying the controller learning rate, color-coded by shade of red). Solid lines represents full model error. Dashed lines correspond to error of the controller memory, obtained by temporarily silencing the adapter during probe trials. **e**, Performance of the distributed learning model in the strong feedback regime. Top: Schematic of the re-adaptation task, in which adaptation is transiently disrupted by a counter perturbation of reversed polarity. Bottom: Performance of the re-adaptation task of the distributed learning model (solid lines) and a plastic controller (dashed). Shade of red represents different consolidation speeds (same legend as in **d**). The distributed learning model shows savings in the re-adaptation phase (i.e., faster re-learning of the 40° perturbation). The plastic controller does not show savings.

Together, these modifications to the error feedback considerably improved the generalization of the controller. The fixed pseudoinverse weights led to far better performance in the perturbed task than standard backpropagation-trained weights, especially in the high-gain regime (Figure 2d-f, Supplementary Figure 2). Importantly, this performance is achieved without altering the original memory encoded in the controller’s recurrent weights: the controller immediately recovers its baseline task performance if the perturbation is removed (Figure 2g). Furthermore, the pseudoinverse constraint provides a simple solution to ensure that the error signal is always properly aligned with the output-potent dimensions of the controller dynamics. Going forward, we will simply assume that this feedback structure has been tuned during development.

Thus far, we have assumed that the controller has access to the error of the motor command, allowing it to correct its internal dynamics in real-time without further training. However, it is not clear how such an internal error signal could be obtained, as sensorimotor errors can only be computed in the external reference frame (i.e., in **y**(*t*)). The presence of the nonlinear motor plant makes it impossible to perform this change of coordinates using a simple linear mapping like the pseudoinverse. Consequently, a secondary module, the adapter, will be required to generate the internal feedback signal in the appropriate coordinates. The following section provides a detailed explanation of its role and structure.

### Dual role of the predicted internal error

We next asked how the internal error signal needed to correct the controller’s dynamics could be produced. One possibility is that the motor system may internally perform a change of coordinates from the behavioral space to the motor command space through hard-wired circuitry. However, perturbations in the plant or the environment may alter the relationship between the motor command and hand position [45], so that any change of coordinates must be continuously re-tuned and adapted over the lifetime of the organism. An alternate solution is to introduce a second *adapter* network that learns a prediction of the error signal. This would decouple the system from real-time, exact error feedback, instead providing the controller with a continuously updated estimate of its error.

What information does the adapter need to form such an approximation? The internal error can be reformulated as a function of the controller’s motor command output **c**(*t*) and the target hand trajectory:

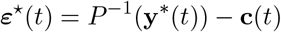

An efference copy of the motor command can directly supply the second term. However, the first term is not necessarily available to the controller, and may even change in response to perturbations of the task (i.e., changes in the target trajectory **y**\*(*t*)). In contrast, because the controller implements an inverse model, the baseline trajectory is already contained in the input **u**(*t*). For small perturbations, the baseline trajectory will be correlated to the new target **y**\*_*pert*_(*t*), and can therefore be used as a proxy for it: 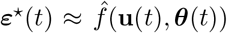 (Figure 3a). In principle, this approximation could be further improved by visual or haptic feedback, supplying information about the mismatch.

This approach not only provides a practical means of estimating the error, but also underpins the functional design of the adapter. Once we have reliable access to both the efference copy and a copy of the upstream input, the adapter no longer needs to learn the full dynamics, and can simply learn a time-invariant function. This can easily be accomplished by multilayer feedforward architectures, known to be universal function approximators in the infinite width limit [46]. Furthermore, in comparison to recurrent circuits, feedforward structures simplify credit assignment during learning and enable more rapid adaptation to sudden changes due to their lack of memory. Structurally, these normative arguments motivate the choice of feedforward structure for the adapter, a choice which is well matched with the cerebellum’s largely feedforward structure [23, 47], open loops with the motor and premotor cortices [48, 49], and roles in fast adaptation [19, 50], and trial-by-trial supervised learning [22].

With the premise that the adapter learns a prediction of the internal error, there are two possible roles for this information. First, the adapter could provide the predicted error signal to the controller, using the same principles of error feedback control investigated in the previous section (Figure 3b, left). Second, the adapter output could bypass the controller entirely, and project directly onto the plant (Figure 3b, center) [41]. Here, the controller and adapter work in parallel as two independent sources of input to the motor plant, with the adapter effectively fine-tuning the baseline motor command produced by the controller.

We tested these possibilities by comparing three different multi-region neural network architectures: *feedback* (the adapter projects to the controller), *parallel* (the adapter projects to the plant) and *hybrid* (the adapter projects to both the controller and the plant; Figure 3b). We fixed the weights of pretrained controller networks and tested each architecture’s ability to adapt to rotations of the target. Adapter weights were trained through backpropagation to minimize the behavioral error, keeping the controller weights fixed (Figure 3c). In the parallel and hybrid architectures we constrained the optimizer in order to prevent the adapter weight updates from using the eligibility trace of the controller state, an important consideration for biological plausibility in multi-region RNNs (this constraint was not possible for the feedback architecture; see Methods). Finally, for comparison, we also tested a single-region plastic controller model.

In the first few hundred epochs, the plastic controller converged faster than the multi-region models (Figure 3c), likely due to the suboptimal initialization of the adapter weights (as the adapter pretraining protocol did not include rotations; see Methods). Nevertheless, both the parallel and the hybrid architectures rapidly achieved comparable performance, even outperforming it at larger learning rates (Figure 3c,d; Supplementary Figure 5). Despite having the fewest constraints in its optimization, the feedback architecture exhibited the poorest performance (Figure 3c,d), consistent with the finding that exact error feedback could only partially generalize to perturbations (cf. the final error in Figure 3c with minimum in 2f). Moreover, direct access to the behavioral output greatly simplifies credit assignment for the adapter module, making the propagation of gradients through the recurrent controller unnecessary (see Methods).

**Figure 5:**
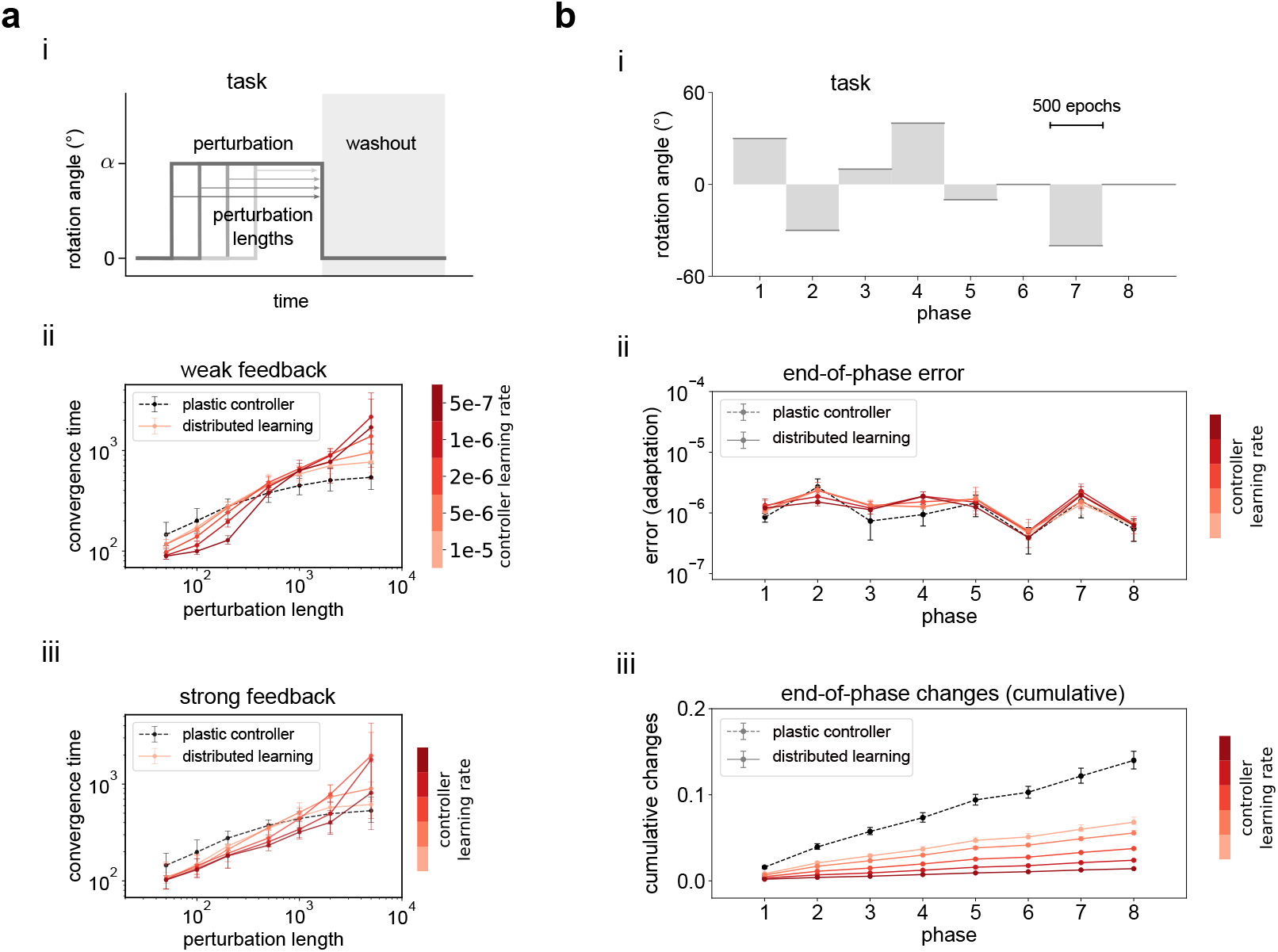
Distributed learning strategy preserves encoded memories in a volatile environment. **a**, Recovery of the baseline memory after transient perturbations. *i*, Schematic of the task. After a transient perturbation, the network must re-learn the baseline memory during a washout phase (grey). *ii*, Speed of re-learning during the washout as a function of the perturbation length (number of epochs required to reach a threshold error value). Shade of red indicates the learning rate in the controller. For comparison, a plastic controller is also shown (dashed black). Feedback gain is *κ* = 1. *iii*, Same as *ii* for high-gain feedback (*κ* = 20). Increasing the gain leads to more robust memory preservation for longer perturbations. **b**, Continual learning task consisting of a sequence of perturbations. *i*, Schematic of the task. The network must adapt to a sequence of random perturbations of varying rotation angle, followed by a final washout phase. *ii*, Final error achieved at the end of each perturbation phase. Different controller learning rates are indicated by the shade of red. A plastic controller network (dashed black) is shown for comparison. All models showed similar performance (solid lines). *iii*, Cumulative changes of the controller’s recurrent weights over different phases. Compared to the plastic controller, distributed learning models resulted in minimal changes of the recurrent dynamics, ensuring preservation of the encoded memory. Color legend is the same for all panels. Error bars indicate min-max values over 5 random seeds.

Together, these results demonstrate that the adapter can be used to approximate the internal error of the controller, and that its access to the plant is crucial for adaptation, both simplifying learning and enabling fine-tuning of the motor command. Indeed, the cerebellum is known to send direct projections to the spinal cord [51], and this projection plays a key role in motor execution and learning [37]. Conversely, because of the similar performance of the parallel and hybrid architectures, it is so far unclear whether the feedback projection to the controller is truly necessary. Motivated by the inter-looped pathways between the cerebellum and motor cortex, we will proceed with the hybrid architecture for the remainder of the manuscript. The question of the feedback projection will be revisited in the final step.

### Distributed learning reproduces slow and fast contributions to adaptation

Thus far, the multi-region networks that we have built have fallen into the controller-adapter hypothesis, in which a fixed controller stores a stable baseline memory and a plastic adapter learns to compensate for perturbations. In the next step, we incorporated slow plasticity in the controller (Figure 4a). Specifically, the controller’s recurrent weights were trained with backpropagation to minimize the error between the behavioral output and the output decoded from the controller’s memory alone (i.e., with the adapter silenced; see Methods). This ensured that the new memory would be fully consolidated in the controller on a timescale determined by its learning rate. As before, the adapter weights were trained to minimize the behavioral error, thereby effectively learning the residual between the target and the controller’s slowly evolving output. Plasticity in the controller and adapter occurred simultaneously, together forming a distributed learning system.

The separation of learning timescales was able to reproduce the two phases of adaptation predicted by classic behavioral studies [5, 6]: a rapid and transient contribution from the fast system (the adapter), and a slow but sustained contribution from the slow system (the controller; Figure 4b). Eventually, the new memory of the perturbed task was fully consolidated in the controller’s intrinsic dynamics, visible in the slow convergence of behavioral trajectories decoded in probe trials with the adapter silenced (Figure 4c). Finally, varying the controller’s learning rate continuously varied the speed at which the new memory was consolidated without compromising the rapid adaptation of the full network performance (Figure 4d; for both weak and strong feedback regimes, Supplementary Figure 6). This supports the idea that the distributed learning strategy enables flexible tuning of the consolidation strategy along the stability-adaptability continuum.

**Figure 6:**
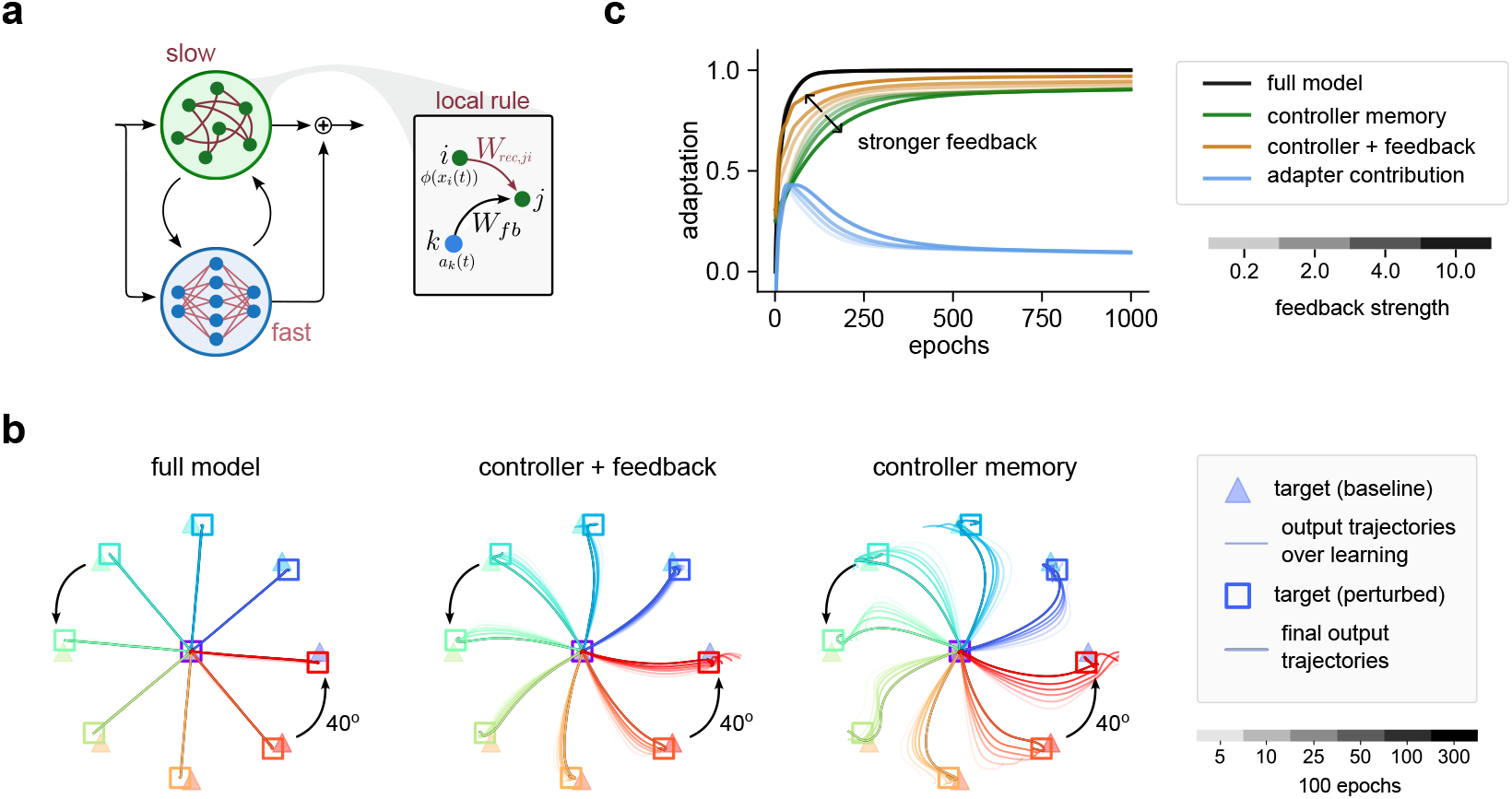
Adapter feedback tutors consolidation of corrected dynamics through a local plasticity rule. **a**, Schematic of the hybrid architecture implementing distributed learning while using a local learning rule in the controller. Inset: the local rule updates each controller recurrent weight proportionally to pre-synaptic activity and its input from the adapter feedback projection. **b**, The evolution of behavioral trajectories during a 40 degree rotation while using local plasticity in the controller (controller local learning rate of 5e-3). Left: the full model rapidly adapts and maintains the correct output. Right: the controller memory slowly changes over the entire course of training, as shown in probe trials in which the adapter is silenced. Center: Behavioral trajectories decoded from the controller, keeping intact the adapter feedback (i.e., only silencing the projection from the adapter to the plant). The adapter feedback drive to the controller plays an important role in correcting the behavioral trajectories throughout learning. Color indicates target direction. Brightness indicates the decoded epoch. **c**, Adaptation level showing module-specific contribution for different gains (feedback strength, indicated by brightness). Stronger feedback increases the contribution of the feedback-driven controller (orange), while slowing the consolidation of the controller intrinsic dynamics (green).

A key advantage of the distributed learning system is the preservation of the encoded memory in the controller. Indeed, multiple timescale learning has been argued to promote faster relearning of a previously encountered perturbation [5], a property known as savings. In order to test for savings in our model, we evaluated relearning of a 40 degree rotation after a brief perturbation of reversed polarity (see Methods; Figure 4e, top). Compared to the plastic controller, our model showed clear evidence of savings consistent with experimental data (particularly in the strong-feedback regime;

Figure 4e, bottom; Supplementary Figure 7). The plastic controller showed faster convergence than the distributed learning system in the initial phase, but slower convergence during relearning. Intuitively, in the distributed learning model, at relearning onset, the controller remained biased towards the original task, reducing the correction that the adapter needed to generate (Supplementary Figure 7b, iii). This is not the case for the plastic controller, since its memory is overwritten during the reversal (Supplementary Figure 7b, iv). This ability to preserve the controller memory may be beneficial in volatile environments where the agent is subject to transient perturbations.

**Figure 7:**
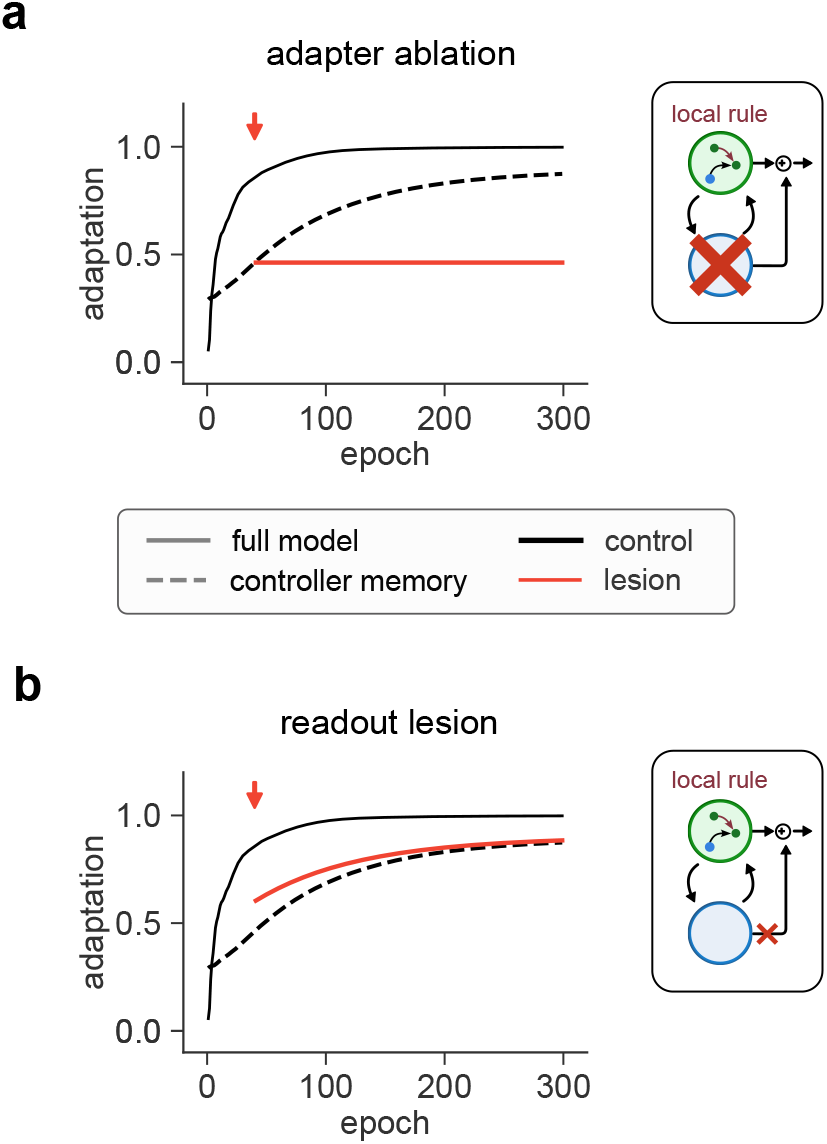
Post-lesion learning as a result of tutored consolidation. Adaptation level for different ablation tests of the full distributed learning model (hybrid architecture, local rule in the controller). For both panels, a specific component of the model was inactivated mid-way through learning (red). For reference, the control (no ablation) is shown in black. Dashed lines indicate consolidated controller memory. **a**, Full ablation of the adapter module. After ablation, performance drops to the level that the controller had consolidated at damage onset. No further learning occurs. **b**, Lesioning the readout projection from the adapter to the plant. Here, the adapter contribution is restricted to its feedback drive and tutoring controller plasticity. After early epochs, the adapter starts encoding an increasingly accurate approximation of the internal error, resulting in continued learning post lesion due to the tutored consolidation mechanism. All simulations used controller local learning rate 5e-3.

To investigate this systematically, we assessed how rapidly the distributed learning system could re-adapt to the baseline task once a transient perturbation is removed (i.e., the washout; Figure 5a, i). Due to the slow consolidation speed, a brief perturbation should lead to little change in the controller, thereby allowing the adapter to rapidly learn the new residual. On the other hand, a sufficiently long perturbation should lead to full consolidation of the updated motor memory in the controller. Indeed, compared to a plastic controller, the distributed learning model more rapidly re-learned the original task for short perturbations, albeit more slowly for long ones (Figure 5a, ii). The ability of the distributed learning model to preserve memories proved even more robust in the strong feedback regime (Figure 5a, iii vs. ii), where it is outperformed by the plastic controller only for extremely long perturbations. These results underscore how the stability–adaptability trade-off depends on environmental volatility, and demonstrate that distributed learning remains effective in most of the tested cases.

In the distributed learning model, each perturbation engenders an early, transient contribution from the adapter that is only gradually offloaded to the controller (Figure 4b), thereby protecting encoded memories from unnecessary modifications if the perturbation is quickly removed. Limiting changes to the recurrent weights could mitigate interference, and therefore forgetting, of the information they encode. We therefore investigated the ability of our model to preserve memories in a continual learning task by subjecting it to a series of perturbations of random strength (Figure 5b, i). The overall performance of the network at the end of each phase was comparable to the plastic controller case (Figure 5b, ii; Supplementary Figure 8). However, the magnitude of changes on the controller recurrent weights was drastically reduced in the distributed learning model, and this amount could be tuned by the controller learning rate (Figure 5b, iii). Thus, we conclude that the distributed learning system enables the preservation of synaptic memories without compromising performance in adaptation and learning.

### Tutored consolidation through local plasticity

Thus far, we have not fully leveraged the feedback projection from the adapter to the controller. When using gradient descent in both regions, each is able to learn independently of the other. Since the adapter learns a prediction of the controller’s internal error, we speculated that the adapter’s feedback projection could be used as a teaching signal for the controller, eliminating its reliance on backpropagation. For this, we reformulated a local error-modulated plasticity rule recently studied in feedback RNNs [42–44], replacing the error signal with the adapter’s approximation already available through the feedback projection. In brief, each recurrent weight was potentiated proportionally to the coactivation of the presynaptic neuron and the adapter projection to the postsynaptic neuron (Figure 6a; see Methods). In this final model, gradient descent is limited to the adapter, justified by classic hypotheses of supervised learning in the cerebellum and unsupervised learning the motor cortex [13].

Just as in the fully backpropagation-trained model, the local rule resulted in two phases of adaptation, with the speed of consolidation determined by its learning rate (Supplementary Figure 9a). However, although the controller memory exhibited the expected convergence toward the rotated targets, it did not achieve the same performance levels as the backpropagation-based consolidation rule (cf. Figure 6b, right and Figure 4c, right). The adapter’s contribution was never entirely suppressed (Supplementary Figure 9b), likely due to the sub-optimality of the local learning rule. Instead, the adapter feedback continued to correct the controller’s dynamics as an external drive, even in later training epochs (Figure 6b, center vs. right). The full system maintained high precision, primarily due to the adapter’s direct projection to the plant (Figure 6b, left vs. center).

The relative importance of the adapter feedback as an external drive versus teaching signal could be modulated through the feedback gain. Increasing the gain led to slower consolidation and therefore a more prolonged contribution from the adapter (Figure 6c, green vs. blue lines). Larger gains also resulted in more effective contribution of the adapter drive to the controller, leading to faster convergence of the output decoded from the feedback-driven controller (Figure 6c, gold lines; Supplementary Figure 9d).We speculate that this effect may be due to the fact that the high-gain regime allowed the driven controller dynamics to more rapidly correct its output, leading to a smaller error predicted by the adapter, and thus a smaller teaching signal for consolidation. High-gain feedback also improved memory preservation in the re-adaptation and continual learning tasks (Supplementary Figures 10-12). Overall, the tradeoff between adapter-driven dynamics and consolidation can be further fine-tuned through gain modulation of the feedback projection, a role which could be implemented in the motor thalamus [52].

Importantly, with the local rule the adapter feedback plays an active role in tutoring consol-idation in the controller. This role can lead to surprising effects in ablation-like experiments in which specific components of the model are inactivated at different stages of learning. Ablating the entire adapter reduced performance to the level of the controller contribution and prevented further adaptation (Figure 7a), with the strongest effect in early ablations before any consolidation could occur (Supplementary Figure 13a). A similar initial drop in performance was observed when the adapter’s projection to the plant was lesioned; surprisingly, however, learning continued even though the adapter itself was no longer able to update its weights (Figure 7b). This paradoxical post-lesion learning occurred only when the adapter’s output was sufficiently aligned with the internal error, allowing the intact feedback projection to keep tutoring the controller (Supplementary Figure 13b). A similar effect on consolidation was observed when freezing the adapter weights mid-way through learning (Supplementary Figure 13c). Finally, lesioning the adapter’s feedback projection prevented consolidation without impacting overall performance (Supplementary Figure 13d). Collectively, these inactivation studies provide clear indicators of the mechanisms underlying our distributed learning model, and clarify the three distinct roles played by the adapter’s predicted error signal: fine-tuning the controller’s output, driving its dynamics via error feedback, and tutoring consolidation of the new memory in the controller.

## Discussion

Flexible motor behaviour requires a balance between stability and adaptability of motor representations. We have advocated for a distributed, two-timescale learning strategy that can tune the balance of this trade-off through the speed at which adapted neural dynamics are consolidated into the original memory. Towards this end, we proposed a multi-region recurrent neural network model composed of a recurrent controller that stores stable memories that are slowly updated over learning, and a feedforward adapter that rapidly learns to respond to perturbations in the environment. Consolidation in the controller occurs simultaneously with adaptation, without requiring separate learning phases or replay, and can be tuned directly through its specific learning rate parameter, or indirectly through the gain of the adapter feedback. This enables the full system to rapidly adapt to external perturbations while controlling the rate of change of the controller recurrent weights. Finally, our work provides a mechanism by which encoded motor memories can be slowly and continuously updated without a concomitant loss in performance.

Our model builds on a large body of work using trained RNNs as generators of motor primitives [43, 53–56], while making specific architectural choices essential to the function of a distributed learning system. In particular, our incorporation of a nonlinear motor plant introduces a mismatch between the external sensorimotor coordinates where errors are computed and the internal coordinates needed to correct the controller [4]. This mismatch motivates the need for an adapter network to produce a prediction of the controller’s internal error. The adapter’s role as a function approximator of the internal error justifies the choice of its feedforward structure, which also simplifies learning due to the lack of recurrent interactions and ensures rapid adaptation. This predicted error signal is then used in three ways. First, it corrects the controller’s internal dynamics using principles of feedback control [42, 44, 55, 56]. Second, it fine-tunes the controller’s output through a direct projection to the plant. Third, the predictive error signal is used to tutor consolidation of the adapted memory using a local plasticity rule. Finally, a crucial component for coordinated learning between the two regions is the fixed pseudoinverse feedback weights, which ensures that the adapter and controller are aligned throughout learning. Plasticity is therefore confined to functionally necessary components (i.e., the internal connectivity in each region), while structured, fixed weights enable effective communication between the controller and the adapter. These hardwired pathways can be interpreted as developmental priors that do not need to be relearned over the animal’s lifetime.

Our distributed learning model reproduces a key prediction from classic behavioral studies positing the existence of interacting slow and fast learning systems [5]. The determinants of the slow and fast systems remains a subject of debate, with alternate proposals including explicit versus implicit learning strategies [6, 12], or differences in contextual inference [57], cell type [58], synapse type [44], or plasticity timescale [59]. In our model, the key factor causing slow and fast learning in the controller and adapter are the speed and efficacy of their respective plasticity rules. Beyond the motor domain, a similar separation of learning timescales has been proposed for episodic memory circuits involving the cortex and the hippocampus [9, 10]. Slow-fast learning has also been argued to underlie habit formation, although here the motor cortex is thought to play the role of the fast learner, with slow learning in the basal ganglia [7, 8]. The identities of the slow and fast learner are likely to be diffuse in the brain, involving subcortical regions which may serve as a final site of stored motor memories [60]. A diversity of learning timescales may thus be a general motif of flexible learning, incorporated at multiple spatial scales and across cognitive and motor behaviors. Based on the functional and structural requirements of our model, we proposed the motor cortex and cerebellum as prime candidates for the controller and adapter. Indeed, the motor cortex has been shown to store stable motor memories over long periods [18, 61, 62], and is thought to implement a dynamical system to control motor output using its recurrent connectivity [15, 63– 66]. In comparison, the largely feedforward structure of the cerebellum [32, 33] has traditionally been linked to learning associations for motor adaptation [20, 23, 67–69]. The cerebellum is also known for its rapid plasticity: a single spike from a climbing fiber (thought to be a teaching signal conveying sensorimotor errors [70, 71]) is sufficient to induce behavioral changes in a single trial [19]. Our model’s separation of learning rules, with backpropagation relegated to the adapter and local plasticity in the controller, are also consistent with the broader roles of supervised and unsupervised learning in the cerebellum and motor cortex, respectively [13, 22]. The interconnected structure between the adapter and controller mirrors the disynaptic pathways forming a parallel organization of cortico-cerebellar loops [25, 49, 72, 73]. The attribution of slow and fast systems to the motor cortex and cerebellum, respectively, is also supported by a transcranial stimulation study in humans, which dissociated rapid cerebellar acquisition of a visuomotor rotation from longterm memory retention in the motor cortex [27]. Finally, while we focused on how learning can be distributed across the cerebellum and motor cortex, such adaptation likely co-exists with upstream changes [3, 74], especially in contexts involving more cognitive strategies [75].

Our work integrates recent developments in RNN dynamics with classic principles from control theory. In particular, our model is a neural network implementation of an inverse model: that is, a system that aims to produce the signal needed to control a fixed plant to achieve a desired motor output [4, 36]. This makes our model functionally distinct from other multi-region models in which a secondary structure is tasked with prediction of future gradients of the cortex [76], performing a low-rank perturbation of cortical dynamics [77], or providing further training to the cortex through offline replay [10]. In particular, a recent modeling study proposed that the cerebellum may drive the cortex with a prediction of its future output [31], essentially implementing a forward model [34, 45]. Both forward and inverse models have been attributed to the cerebellum, working in combination in paired circuits [45]. Our work refines this view, by forming an inverse model with both controller and adapter together, an assumption supported by recent evidence of motor cortical activity encoding an inverse dynamics model of the musculoskeletal model system [78].

Another distinctive feature of our model is the inclusion of a direct projection from the adapter to the plant, bypassing the controller. The adapter-plant projection simplifies the adapter’s credit assignment problem during learning, enabling rapid adaptation to environmental changes without access to the controller’s hidden unit activity. This architectural choice enables flexibility in the specific form of the adapter learning rule: as long as the adapter learns a good approximation of the residual needed to drive the plant, that same error signal can begin correcting and consolidating the controller’s dynamics using the local rule. Surprisingly, the local rule did not necessarily decrease performance when compared to backpropagation-based consolidation (Supplementary Figure 12), suggesting that suboptimal learning in the controller can be compensated by the adapter. This would not be possible with the common feedback-only architecture used in many multi-region models [31, 53, 76, 77], and opens the possibility of using more reinforcement-like learning rules [79–81], though trial-and-error perturbations may substantially slow adaptation. Clarifying the origin of the teaching signal is a key challenge for future research.

Our hypothesis that the adapter tutors consolidation in the controller yields a number of experimental predictions for cerebellar inactivation studies during motor adaptation. A common prediction is that late inactivations should have small effect, as the new memory will already be consolidated in the controller. In line with this prediction, silencing the majority of cerebellar neurons has been shown to compromise learning, without strongly affecting normal motor behavior [82]. Our model also predicts that cerebellar learning should persist even after behavioral performance is optimized, to compensate for slow changes in the controller memory throughout consolidation.

Indeed, studies of short-term saccadic adaptation have shown that Purkinje cell complex spike activity, the primary teaching signal in the cerebellum, remains high even after behavioral errors are minimized [83], possibly reflecting a cerebellar role in ongoing consolidation. Our model of tutored consolidation is also supported by recent evidence showing that inactivating the cerebello-spinal projection affects early stages of motor learning in a rotarod task, while having minimal effect at late stages [37]. Intriguingly, silencing contralaterally-projecting cerebellospinal neurons during early stages of the rotarod task only partially impaired learning, as performance continued to increase at a diminished rate [37]. This mirrors the paradoxical post-lesion learning that we observed when silencing the adapter readout projection. However, compensatory plasticity mechanisms in other brain regions could also account for this continued learning, as the rotarod task is known to engage striatal circuits [84]. Targeted inactivation studies in more constrained, cerebellar-dependent tasks will be essential to directly test the tutored consolidation mechanism proposed here.

The environment never stops changing, forcing organisms to learn continuously throughout their lives. Recent interest in sequential and continual learning has has led to a variety of strategies aimed to avoid catastrophic forgetting [40, 85], such as encoding task-specific dynamics in orthogonal latent spaces [86]. However, these solutions can be inefficient when the new task is a variation of previous task, since it may fail to support generalisation. In such cases, a two-timescale learning strategy can preserve encoded memories while acting in a volatile environment. This may also be useful for tasks that change continuously over the lifetime of the animal, where indexing by discrete context [57, 87] becomes inefficient. A combination of these approaches is likely to be employed for flexible motor learning. Understanding how the brain distinguishes between adaptable tasks and novel skills will be a key question for future work.

## Methods

### Network dynamics and architectures

#### Controller network

The controller is defined as a continuous-time recurrent neural network (RNN) obeying the following equation:

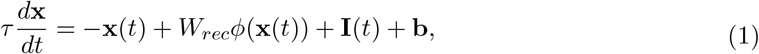

where **x**(*t*) ∈ ℝ^*n*^ represents the hidden unit states, *W*_*rec*_ ∈ ℝ^*n×n*^ are the recurrent weights, *ϕ* is a nonlinear activation function (tanh), **b** ∈ ℝ^*n*^ are unit-wise biases, and *τ >* 0 is the membrane time constant. The total input **I**(*t*) is the sum of two input sources:

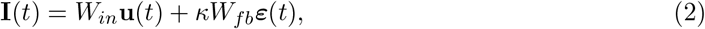

where **u**(*t*) ∈ ℝ^*m*^ are external inputs with weights *W*_*in*_ ∈ ℝ^*n×m*^, and ***ε***(*t*) ∈ ℝ^*d*^ is a feedback signal with weights *W*_*fb*_ ∈ ℝ^*n×d*^. Depending on the architecture, ***ε***(*t*) will either represent the exact internal error or an approximation learned by the adapter (see ‘Feedback’ below, Table 1). The free parameter *κ* ≥ 0 represents the strength of the feedback (*feedback gain*; unless otherwise specified, we set *κ* = 1). The output **c**(*t*) ∈ ℝ^*d*^ of the controller is defined as:

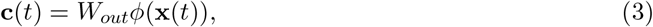

**Table 1.**
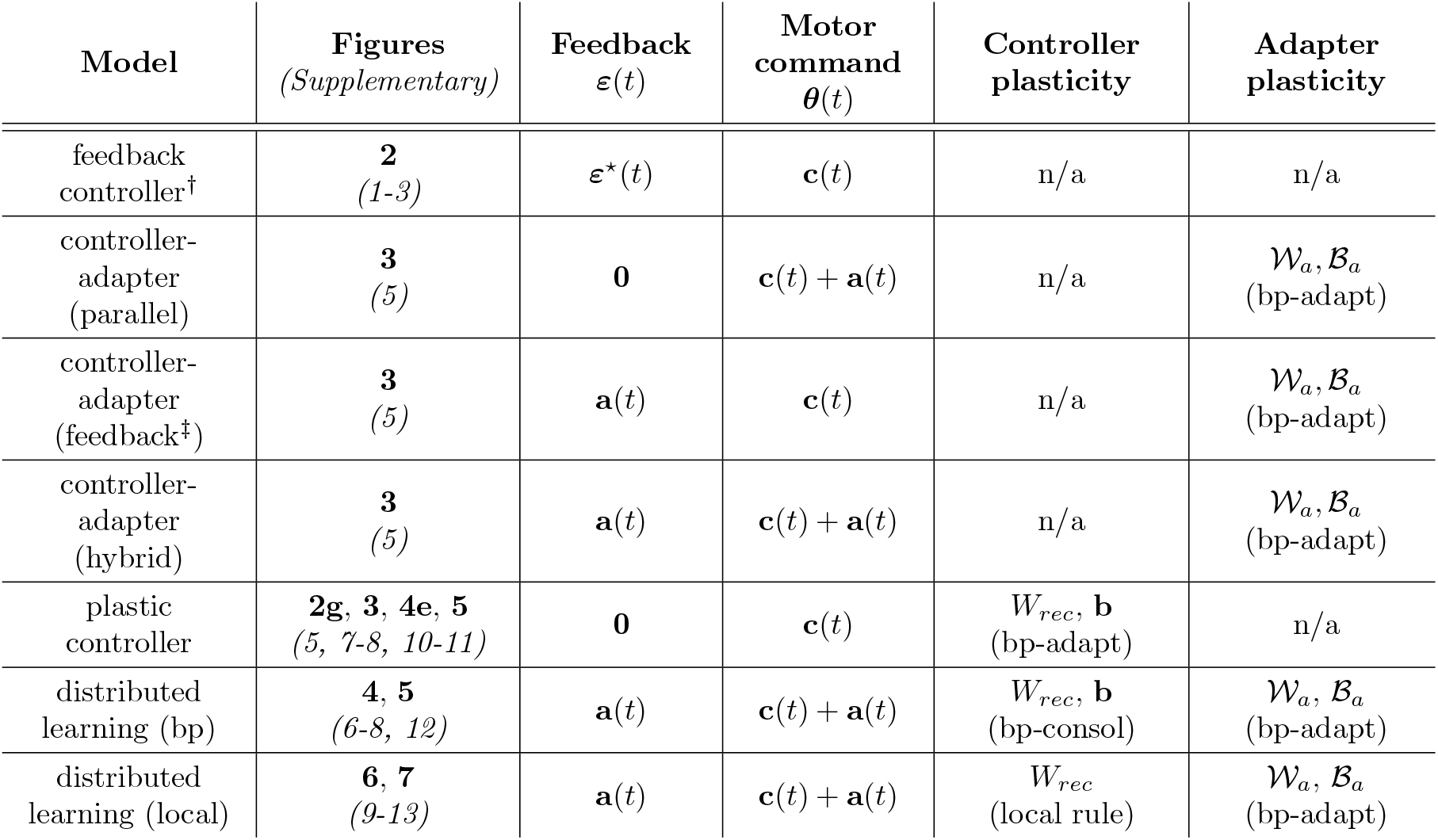
Architecture and plasticity of all models. ^†^ The feedback controller is the only model that integrates real-time sensorimotor error feedback. ^‡^ The controller-adapter (feedback architecture) is the only model in which the adapter weight updates used shared gradients (see ‘Backprop for adaptation (bp-adapt)’).

| Model | Figures<br>(*Supplementary*) | Feedback<br>$\varepsilon(t)$ | Motor<br>command<br>$\theta(t)$ | Controller<br>plasticity | Adapter<br>plasticity |
| --- | --- | --- | --- | --- | --- |
| feedback<br>controller<sup>†</sup> | **2**<br>(1-3) | $\varepsilon^*(t)$ | $\mathbf{c}(t)$ | n/a | n/a |
| controller-adapter<br>(parallel) | **3**<br>(5) | **0** | $\mathbf{c}(t) + \mathbf{a}(t)$ | n/a | $\mathcal{W}_a, \mathcal{B}_a$<br>(bp-adapt) |
| controller-adapter<br>(feedback<sup>‡</sup>) | **3**<br>(5) | $\mathbf{a}(t)$ | $\mathbf{c}(t)$ | n/a | $\mathcal{W}_a, \mathcal{B}_a$<br>(bp-adapt) |
| controller-adapter<br>(hybrid) | **3**<br>(5) | $\mathbf{a}(t)$ | $\mathbf{c}(t) + \mathbf{a}(t)$ | n/a | $\mathcal{W}_a, \mathcal{B}_a$<br>(bp-adapt) |
| plastic<br>controller | **2g, 3, 4e, 5**<br>(5, 7-8, 10-11) | **0** | $\mathbf{c}(t)$ | $W_{rec}, \mathbf{b}$<br>(bp-adapt) | n/a |
| distributed<br>learning (bp) | **4, 5**<br>(6-8, 12) | $\mathbf{a}(t)$ | $\mathbf{c}(t) + \mathbf{a}(t)$ | $W_{rec}, \mathbf{b}$<br>(bp-consol) | $\mathcal{W}_a, \mathcal{B}_a$<br>(bp-adapt) |
| distributed<br>learning (local) | **6, 7**<br>(9-13) | $\mathbf{a}(t)$ | $\mathbf{c}(t) + \mathbf{a}(t)$ | $W_{rec}$<br>(local rule) | $\mathcal{W}_a, \mathcal{B}_a$<br>(bp-adapt) |

**Table 2.** Main symbols used in the paper.

| Symbol | Definition |
| --- | --- |
| $\phi(\cdot)$ | Recurrent module activation function (tanh) |
| $\psi(\cdot)$ | Adapter module activation function (ReLU) |
| $W_{in}$ | Input weights, controller |
| $W_{rec}$ | Recurrent weights, controller |
| $W_{out}$ | Output weights, controller |
| $W_{fb}$ | Feedback weights, controller |
| $P(\cdot)$ | Motor plant |
| $\kappa$ | Feedback gain |
| $\mathbf{u}(t)$ | External input |
| $\mathbf{x}(t)$ | Hidden layer internal state |
| $\boldsymbol{\varepsilon}^*(t)$ | Internal error |
| $\boldsymbol{\varepsilon}(t)$ | Feedback to the network |
| $\mathbf{c}(t)$ | Controller output |
| $\mathbf{a}(t)$ | Adapter output |
| $\boldsymbol{\theta}(t)$ | Motor command |
| $\mathbf{y}(t)$ | Behavioral output |
| $\cdot^*$ | Target values |
| $\tilde{\mathbf{y}}(t)$ | Controller memory output |
| $\tilde{\tilde{\mathbf{y}}}(t)$ | Controller+feedback output |

where *W*_*out*_ ∈ ℝ^*d×n*^.

#### Plant and motor comman

The controller output generates behavioral trajectories **y**(*t*) ∈ ℝ^*l*^ indirectly by providing input to the plant, here defined as a nonlinear, invertible function *P* : ℝ^*d*^→ ℝ^*l*^, which maps motor command signals to behavioral output: **y**(*t*) = *P* (***θ***(*t*)). Depending on the architecture of the model, the motor command ***θ***(*t*) ∈ ℝ^*d*^ is either defined as the controller output, or the combined output of the controller and the adapter:

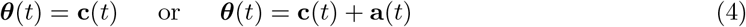

where **a**(*t*) ∈ ℝ^*d*^ is the adapter output (see ‘Adapter network’ below, Table 1).

#### Internal erro

For any target behavioral trajectory **y**\*(*t*), we can define the target motor command by inverting the plant nonlinearity:

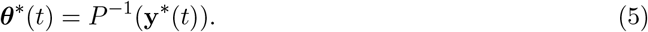

This quantity can then be used to define the controller’s *internal error* :

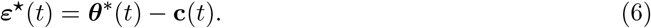

i.e., the difference between the target motor command and the controller output.

#### Feedback

Depending on the architecture of the model, the feedback ***ε***(*t*) to the controller may take one of two forms:

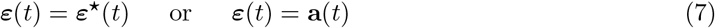

In the first case, the feedback is the real-time internal error (also referred to as *exact error feedback*). This allows the controller to adjust its dynamics for online correction following principles of feedback control. In the second case (used for all multi-area models, Table 1), the feedback is equal to the adapter output. In principle, if ***θ***(*t*) = **c**(*t*) + **a**(*t*) (i.e., in the hybrid architecture), and if **c**(*t*) is stationary over trials (either because the controller weights are fixed, or because controller plasticity is much slower than adapter plasticity), ***ε***⋆(*t*) can be considered as the target adapter output: **a**\*(*t*) = ***ε***⋆(*t*). For this reason, the adapter feedback is also referred to as the *predicted error*.

Unless otherwise specified, the feedback weights are set to the Moore-Penrose pseudoinverse of the output weights of the controller module:

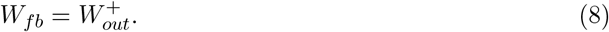

The only exceptions are in Figures 2f and Supplementary Figures 1-2, where *W*_*fb*_ is optimized alongside other parameters during pretraining (referred to as *trained* feedback weights) for comparison to the pseudoinverse solution.

#### Adapter network

The adapter is defined as a two-layer feedforward network with *n*_*a*_ units per layer, and activation function *ψ* (ReLU). It receives a copy of both the external inputs **u**(*t*) and the controller’s output **c**(*t*). The adapter output **a**(*t*) ∈ ℝ^*d*^ is generated by a third, linear readout layer. The adapter network weights and biases are denoted by the sets 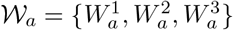, where 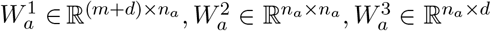 and 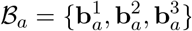, where 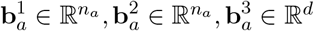, respectively.

#### Controller memo

In the distributed learning models (Table 1), we quantified the amount of consolidation in the controller by testing its performance with the adapter silenced. Specifically, we first simulated the controller output in the absence of feedback:

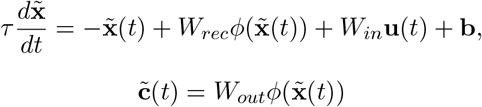

The behavioral output in this ‘no-adapter’ case is then given by 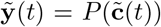. In Figure 6 and Supplementary Figure 9, we also quantify the feedback-driven memory, i.e. the one obtained when removing its direct contribution to the plant while keeping the adapter’s projection to the con-troller. In this case, we obtain 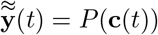 (‘controller+adapter feedback’).

#### Overview of tested models

The various models tested in our analyses differed via architecture and/or plasticity sites. Architectural differences were determined by the source of the error feedback, and the definition of the motor command. These correspond to major differences in the interconnectivity between the controller, adapter, and plant. Differences in plasticity were determined by which parameters were plastic during adaptation or by how those parameters evolved. All variations are summarized in Table 1. For all models, dynamics were simulated using the Euler method with step size *dt* (see Table 3).

**Table 3.** Parameters and variables. G.D. stands for *Gradient Descent*. Empty values indicates parameters that are systematically varied (i.e., learning rates).

| Parameter | Definition | Value |
| --- | --- | --- |
| $m$ | N. inputs | 3 |
| $d$ | N. motor command dimensions | 2 |
| $n$ | Controller hidden units | 256 |
| $n_a$ | Adapter hidden units | 1024 |
| $dt$ | Time step | 10 ms |
| $\tau$ | Recurrent module time constant | 100 ms |
| $T$ | Total movement time | 1300 ms |
| $T_{no-go}$ | Rest time before movement | 300 ms |
| $T_{mov}$ | Movement time | 1000 ms |
| $B$ | N. reaching trajectories (equiv. batch size) | 8 + 1 |
| $r$ | Endpoints circle radius | 10 cm |
| $v$ | Speed constant | 10 m/s |
| $l_1$ | Arm length | 22 cm |
| $l_2$ | Forearm length | 27 cm |
| $s_1, s_2$ | Shoulder position | (0cm, 0cm) |
| $B_{pre,a}$ | Adapter pretraining batch size | 3500 |
| $N_{epochs, pre, c}$ | G.D.: controller pretraining epochs | 50000 |
| $N_{epochs, pre, a}$ | G.D.: adapter pretraining epochs | 20000 |
| $N_{reg drop}$ | G.D.: regularizers drop epoch | 5000 |
| $\eta_{pre, c}$ | G.D.: learning rate, controller, pretraining | 1e-4 |
| $\eta_{pre, a}$ | G.D.: learning rate, adapter, pretraining | 1e-4 |
| $\eta_c$ | G.D.: learning rate, controller, adaptation | - |
| $\eta_a$ | G.D.: learning rate, adapter, adaptation | - |
| $\eta_{local}$ | G.D.: learning rate, controller, adaptation, local rule | - |
| $\nu$ | G.D.: L-2 weight regularisation | 1e-4 |
| $\gamma_{in}, \gamma_{rec}, \gamma_{out}$ | nonlin. gain factors | Pytorch default values |
| $\beta_1, \beta_2$ | Beta parameters for adam optimizer | (0.9, 0.999) |

#### Network plasticity

Here, we describe the three forms of plasticity used to train models during all adaptation tasks. This does not include parameter pretraining, which was used to initialize parameters in the controller and adapter prior to adaptation (see ‘Pretraining protocols’ section below). Region-specific plastic parameters are shown in Table 1 for each tested model. All trainings were run in full batch mode, with updates averaged over all *B* = 9 task conditions (corresponding to 8 reach directions plus a no-movement condition). In the consolidation rules (bp-consol and local rule), the corresponding average over each condition *b* was omitted for notational clarity.

#### Backprop for adaptation (bp-adapt)

Here, plastic parameters were optimized to minimize the mean squared error (MSE) of the behavioral trajectory, using backpropagation through time:

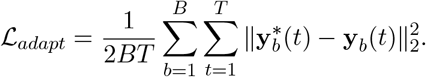

Here, the index *b* indicates task condition (i.e., reach direction; generally dropped for clarity). When present, the adapter was always trained with bp-adapt (Table 1). Unless otherwise specified, optimization was always performed while preventing gradient sharing between the two modules. Specifically, for each module, the output of the other module was detached from the computational graph determining the backpropagated errors - effectively blocking credit assignment through the opposing module (for the adapter, this reduces optimization to simple backpropagation, i.e., not over time, since there’s no recurrence left in the computational graph). The sole exceptions are the feedback controller-adapter architecture (for which using shared gradients is required for training), and Supplementary Figure 5 (where the shared gradients in the hybrid architecture are tested for validation and comparison).

#### Backprop for consolidation (bp-consol)

As a step towards the full model, we tested the ability of the controller to consolidate new memories in the distributed learning architecture using backpropagation through time to minimize the loss function:

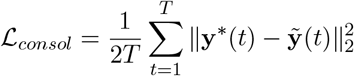

where 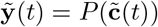 is the behavioral output decoded from the controller with the adapter silenced (see ‘Network dynamics and architectures – Controller memory’ above).

#### Local rule for consolidation

In the local distributed learning model, bp-consol in the controller was replaced with a local plasticity rule. Here, the recurrent weight from neuron *i* to *j* was updated using the following plasticity rule:

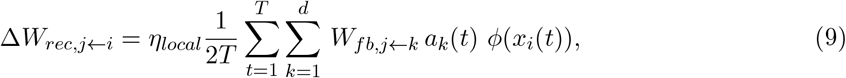

i.e., each recurrent weight in the controller was updated proportional to co-activation of the presynaptic neuron with its predicted error signal input from the adapter. The feedback gain *κ* parameter was not included in the local rule to avoid changing the effective learning rate.

#### Task details

##### Motor plant

The plant *P* is modeled as a static, two-link planar mapping from the shoulder and elbow angles, given by the motor command ***θ***(*t*), to the corresponding hand position:

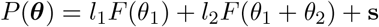

where *F* : ℝ → ℝ^2^ with *F* (*x*) = [cos(*x*), sin(*x*)]^⊤^, **s** ∈ ℝ^2^ represents the shoulder position, and *l*_1_, *l*_2_ *>* 0 represent the lengths of the upper arm and forearm, respectively.

##### Baseline center-out task

For all models, the controller was initially trained to reproduce a set of planar center-out reaching trajectories (details in ‘Pretraining protocols’). The task consisted of *B* = 9 different conditions (8 reach directions plus a no-movement condition). The trajectory for each condition (except no-movement) consisted of an initial no-go period *T*_*no*−*go*_, followed by a movement period *T*_*mov*_ during which the hand position followed a linear trajectory toward one of the eight endpoints *Y*_*b*_ ∈ ℝ^2^, arranged on a circle with radius *r*. The speed along each target trajectory followed a sigmoidal profile, resulting in:

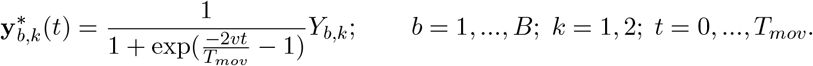

Here, *v* represents the speed constant, and k is the index over the two cartesian coordinates. The input consisted of the reference trajectory and a no-go cue: 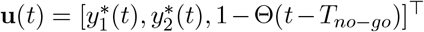, where Θ is the Heaviside function. With this input, the neural network must generate the motor command to drive the plant along the specified trajectories. This makes the network learn an inverse model of the plant.

##### Perturbed task

After pretraining, the network was exposed to a set of perturbations applied to the initial center-out reaching task. While the inputs remained unchanged, the target trajectories were modified by applying different transformations to the initial targets:

- rotations: 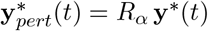, with 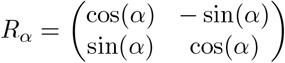;
- translations: 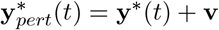, with **v** the translation vector;
- radius changes: 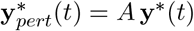, with *A* a constant.

Throughout the main text, we focus on rotations; translations and radius changes are studied only in the feedback controller model, as shown in Supplementary Figure 2. Unless otherwise specified, an *α* = 40° counterclockwise rotation was used (*standard adaptation task*).

##### Equivalence to visuomotor rotations

In our formulation of the perturbed task, the target **y**\*(*t*) is modified while leaving unchanged the mapping from ***θ***(*t*) to **y**(*t*). Here, we note that this is equivalent to a perturbation in the mapping while leaving the target unchanged, e.g., in visuomotor rotations commonly studied in motor adaptation experiments [3]. Specifically, a visuomotor rotation corresponds to adding a linear transformation between the motor command and behavioral output: **y**(*t*) = *R*_*α*_*P* (***θ***(*t*)), where *R*_*α*_ is the rotation matrix:

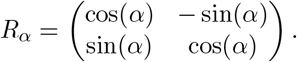

To reproduce the desired behavioural output **y**\*(*t*) after applying the visuomotor rotation, the (updated) motor command must take the form 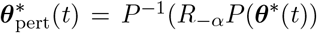, where ***θ***\*(*t*) is the target motor command of the baseline task. Indeed:

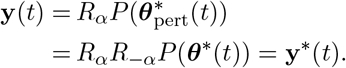

If the perturbation is not applied, the same motor command 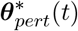 leads to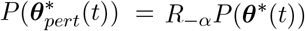, i.e., the behavioral target output rotated by −*α* degrees. Thus, the two formulations are equivalent. A similar argument applies to translations or changes in the radius of the target trajectories.

##### Pretraining protocols

For all analyses, simulations were conducted over 5 random seeds. All models and trainings were implemented in Pytorch [88].

##### Controller pretraining

To obtain stereotypical trajectory dynamics, we first pretrained the controller module to perform the baseline center-out reaching task. All trained parameters were inJitialized using He uniform initialization [89], i.e. from the distribution 𝒰 [−*p, p*], where 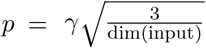 for weights and 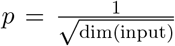 for biases. Models were optimized through single-batch training using the loss function

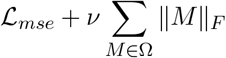

where ℒ_*mse*_ is the mean-squared error over the output trajectory (equivalent to ℒ_*adapt*_), *ν* is a regularization coefficient, ∥ · ∥_*F*_ is the Frobenius norm, and Ω is the set of all trained parameters. Two versions of the model were trained depending on the structure of the feedback: trained feedback weights, and pseudoinverse feedback weights. In the former case, the trained weights were Ω = {*W*_*in*_, *W*_*rec*_, *W*_*out*_, *W*_*fb*_, **b**}. In the pseudoinverse case, Ω = {*W*_*in*_, *W*_*rec*_, **b**}, while fixing 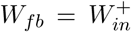. In both cases, parameters were trained for *N*_*epochs, pre, c*_ epochs using Adam optimization with an initial learning rate *η*_*pre, c*_, while applying a cosine annealing down to 0.01*η*_*pre, c*_ by the final epoch. The regularization constant *ν* was also adaptive, and reduced to 0.01*ν* after *N*_*reg drop*_ epochs. In the case of trained feedback weights, similarly to [43], these were included in the optimization only after *N*_*fb,start*_ epochs.

##### Adapter pretraining

We pretrained the adapter module to approximate the internal error ***ε***⋆(*t*) under non-rotational perturbations. Starting from the baseline task, we applied translations (sampled values on a square grid ranging from −2cm to 2cm, with steps of 1cm) and radial scaling (trajectories rescaled by a factor ranging from 0.85 to 1.15, 0.05 steps). The model was trained to approximate the residuals between baseline and perturbed motor commands over time. Specifically, given the baseline motor command ***θ***_*base*_(*t*) and the perturbed motor command ***θ***_*pert*_(*t*) (defined from the specific target trajectories), the adapter output **a**(*t*) was trained to minimize the mismatch between the current motor command ***θ***(*t*) = **c**(*t*) + **a**(*t*) and the target motor command ***θ***\*(*t*) in both cases:

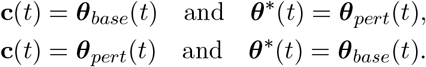

The loss was computed as the MSE between **c**(*t*) + **a**(*t*) and ***θ***\*(*t*), and the set of trained parameters was always Ω = 𝒲_*a*_ ∪ ℬ_*a*_. Weights in 𝒲_*a*_ were initJialized with the Xavier normal initialization [90], using values drawn from 𝒩 (0, *σ*^2^), with 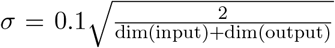 defined over the specific input/output layer dimensions. Biases in ℬ_*a*_ were initialized to zero. Training was run over *N*_*epochs, pre, a*_ epochs, with batch size *B*_*pre, a*_, using Adam optimizer with learning rate *η*_*pre, a*_. A scheduler (`ReduceLROnPlateau` implemented in PyTorch) was used to adaptively reduce the learning rate when the loss plateaued. Results were averaged over 5 random seeds.

##### Adaptation training protocols

For all analyses, simulations were conducted over the 5 different pretrained models, obtained as described in ‘Pretraining protocols’. In the plastic controller case, a standard RNN was used (no feedback). All models and training procedures were implemented using Pytorch [88]. Bp-adapt optimization (see ‘Network plasticity - Backprop for adaptation’) was used for the plastic controller model and the adapter in both controller-adapter and distributed learning models), while either bp-consol or the local rule was used for the controller. For both bp-adapt and bp-consol, Adam optimization was used with default values for the momenta *β*_1_, *β*_2_ (Table 3, [91]). For all adaptation tasks, the learning rate followed a cosine annealing schedule, gradually decreasing to 1% of its initial value, reached at the end of a task-specific time window, for improving convergence. The annealing time window was set to min(5000, *N*_*epochs*_) epochs for the standard adaptation task and ablation tests, and to 2000 epochs for the re-adaptation, washout (washout phase only) and continual learning tasks. The plastic controller’s learning rate was fixed at 1 × 10^−4^, above which the loss curve tended to show difficulties with convergence. The learning rates in bp-adapt for distributed learning models (adapter parameters) were set to either 1 × 10^−4^ or 2 × 10^−4^, depending on the task (see the corresponding task section for details).

##### Standard adaptation task

Networks were trained to minimize the MSE on the perturbation task (40° rotation of the target trajectories, see ‘Task details - Perturbed task’). For all cases, the input signal **u**(*t*) remained unchanged. When training distributed learning (local) models, a consolidation learning rate of 5 × 10^−3^ was used, unless otherwise specified. For bp-adapt (adapter parameters), learning rate was set to 1 × 10^−4^.

##### Re-adaptation task

Models were tested on a 3-phase task, composed by a first 40° rotation, followed by a short counter rotation of the same amplitude lasting 50 epochs, and a final re-adaptation phase, where the initial 40° rotation was restored. For the entire task, inputs were left equivalent to the baseline task. For bp-adapt (adapter parameters), we used a learning rate of 2 × 10^−4^ for the distributed learning models.

##### Washout task

Models were tested on the standard perturbed task, with variable length of perturbation (50, 100, 200, 500, 1000, 2000 or 5000 epochs), followed by a washout period of 5000 epochs (i.e., return to the baseline task). For all models (distributed learning and plastic controller), during washout, learning rate was reset to the initial value and the cosine annealing window for the adapter’s optimization was reduced to 2000 epochs for improving convergence. For bp-adapt, the learning rate (adapter parameters) was set to 1 × 10^−4^.

##### Continual learning task

Models were tested on a sequence of perturbations, each 500 epochs long, of random rotation amplitudes, followed by a final washout phase of 500 epochs. The same sequence was used for all models and seeds. For bp-adapt, the learning rate (adapter parameters) was set to 1 × 10^−4^.

##### Ablation tests

We tested learning in the distributed learning model (local) while inactivating specific pathways at either early (epoch 0), mid (epoch 40), or late (epoch 200) stages. Results were compared to a control with no inactivation. Learning rates were set to 1 × 10^−4^ for bp-adapt (adapter parameters) and 5 × 10^−3^ for the local rule (in the controller). The cosine annealing time window for bp-adapt learning rate was set to 5000 epochs.

##### Analyses

In all plots, error bars or shaded areas are defined as the min-max values among all 5 seeds.

##### Error and adaptation metrics

When plotting errors (either *error (baseline)* or *error (adaptation)*), we refer to the MSE averaged over the full trajectory and across all conditions (as defined in ‘Network plasticity - Backprop for adaptation (bp-adapt)’). Errors are z-scored, using the mean and variance of the corresponding target trajectory. The *adaptation* metric is defined directly from the training error. Let *e*(**y**(*t*)) = ℒ_*mse*_(**y**(*t*)) be the MSE of the trajectory **y**(*t*) with respect to the target **y**\*(*t*). The adaptation metric *ρ* is then:

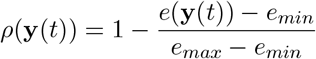

where *e*_*min*_ and *e*_*max*_ are computed numerically, from the full model training error sequences (using control model in the ablation tests; Figure 7, Supplementary Figure 13), over training time and seeds. We define *controller contribution* to be the adaptation metric computed on the behavioral output obtained when the adapter is silenced, i.e.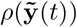. *Controller + feedback contribution* refers to the adaptation metric computed when only the adapter-plant projection is silenced, i.e. 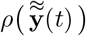. Finally, the *adapter contribution* is defined as the difference between full model adaptation and the controller contribution, i.e. 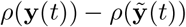.

##### Feedback strength

The total strength of the feedback depends on two factors: the parameter *κ* and the norm of the feedback weights *W*_*fb*_. We defined the *effective feedback strength* as *κ*∥*W*_*fb*_∥_*_, where ∥ · ∥_*_ is the nuclear norm (i.e., the sum of the singular values).

##### Convergence time

In the washout procedure, models were evaluated using a convergence time metric. This was defined as the number of epochs required to reach a threshold error value of 10^−6^.

##### Cumulative recurrent weights changes during continual task

In the continual procedure, we tracked the cumulative changes during training over the recurrent weights matrix *W*_*rec*_, to assess preservation of information in the controller module when the network is subjected to perturbations. Cumulative changes where computed as:

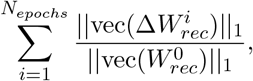

where 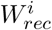 and 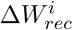 are the recurrent weights and weight updates on the *i*th epoch during training.

## Code availability

Code will be made available on GitHub upon publication.

## Acknowledgements

We thank Joao Barbosa, Angus Chadwick, Matt Getz, Harsha Gurnani, Ashok Litwin-Kumar, Yuxiu Shao, and Heike Stein for their helpful comments on the manuscript. We additionally thank Dhruva Rhaman, Sophie Deneve, and Sara Solla for key feedback on an early stage of the project. This project was supported by the Agence National de Recherche (ANR-20-CE37-0004, ANR-17-EURE-0017, ANR-23-IACL-0008) and INSERM.

## Notes

### Competing Interest Statement

The authors have declared no competing interest.

## References

[1] Reza Shadmehr and Ferdinando A. Mussa-Ivaldi. Adaptive representation of dynamics during learning of a motor task. In Journal of Neuroscience, 1994. URL https://api.semanticscholar.org/CorpusID:6332563.

[2] Sergey D. Stavisky, Jonathan C. Kao, Stephen I. Ryu, and Krishna V. Shenoy. Trial-by-Trial Motor Cortical Correlates of a Rapidly Adapting Visuomotor Internal Model. J. Neurosci., 37(7):1721–1732, 2017. doi: 10.1523/JNEUROSCI.1091-16.2016. URL https://www.jneurosci.org/content/37/7/1721.

[3] Matthew G. Perich, Juan A. Gallego, and Lee E. Miller. A Neural Population Mechanism for Rapid Learning. Neuron, 100(4):964–976.e7, 2018. doi: 10.1016/j.neuron.2018.09.030. URL https://www.cell.com/neuron/abstract/S0896-6273(18)30832-8.

[4] Wolpert Jörn Diedrichsen, and Randall Flanagan. Principles of sensorimotor learning. Nature Reviews Neuroscience, 12:739–751, 2011. URL https://api.semanticscholar.org/ CorpusID:5172329.

[5] Maurice A Smith, Ali Ghazizadeh, and Reza Shadmehr. Interacting adaptive processes with different timescales underlie short-term motor learning. PLoS biology, 4(6):e179, 2006. doi: 10.1371/journal.pbio.0040179.

[6] David M. Huberdeau, John W. Krakauer, and Adrian M. Haith. Dual-process decomposition in human sensorimotor adaptation. Current Opinion in Neurobiology, 33:71–77, 2015. URL https://api.semanticscholar.org/CorpusID:12217322.

[7] F. Gregory Ashby, Benjamin O. Turner, and Jon C. Horvitz. Cortical and basal ganglia contributions to habit learning and automaticity. Trends in Cognitive Sciences, 14(5):208– 215, 2010. doi: 10.1016/j.tics.2010.02.001. URL https://www.sciencedirect.com/science/article/pii/S1364661310000306.

[8] James M. Murray and G. Sean Escola. Remembrance of things practiced with fast and slow learning in cortical and subcortical pathways. Nat Commun, 11(1):1–12, 2020. doi: 10.1038/s41467-020-19788-5. URL https://www.nature.com/articles/s41467-020-19788-5.

[9] James L. McClelland, Bruce L. McNaughton, and Randall C. O’Reilly. Why there are complementary learning systems in the hippocampus and neocortex: insights from the successes and failures of connectionist models of learning and memory. Psychological review, 102 3:419–457, 1995. URL https://api.semanticscholar.org/CorpusID:2832081.

[10] Weinan Sun, Madhu Advani, Nelson Spruston, Andrew Saxe, and James E Fitzgerald. Organizing memories for generalization in complementary learning systems. Nature neuroscience, 26(8):1438–1448, 2023.

[11] Konrad Paul Kording, Joshua B. Tenenbaum, and Reza Shadmehr. The dynamics of memory as a consequence of optimal adaptation to a changing body. Nature Neuroscience, 10:779–786, 2007. URL https://api.semanticscholar.org/CorpusID:13086131.

[12] Jordan A. Taylor, John W. Krakauer, and Richard B. Ivry. Explicit and Implicit Contributions to Learning in a Sensorimotor Adaptation Task. J. Neurosci., 34(8):3023–3032, 2014. doi: 10.1523/JNEUROSCI.3619-13.2014. URL https://www.jneurosci.org/lookup/doi/10.1523/JNEUROSCI.3619-13.2014.

[13] K. Doya. Complementary roles of basal ganglia and cerebellum in learning and motor control. Current Opinion in Neurobiology, 10(6):732–739, 2000. doi: 10.1016/s0959-4388(00)00153-7.

[14] Daniele Caligiore, Giovanni Pezzulo, Gianluca Baldassarre, Andreea C. Bostan, Peter L. Strick, Kenji Doya, Rick C. Helmich, Michiel Dirkx, James Houk, Henrik Jörntell, Angel Lago-Rodriguez, Joseph M. Galea, R. Chris Miall, Traian Popa, Asha Kishore, Paul F. M. J. Verschure, Riccardo Zucca, and Ivan Herreros. Consensus Paper: Towards a Systems-Level View of Cerebellar Function: The Interplay Between Cerebellum, Basal Ganglia, and Cortex. Cerebellum, 16(1):203–229, 2017. doi: 10.1007/s12311-016-0763-3. URL https://link.springer.com/article/10.1007/s12311-016-0763-3.

[15] Saurabh Vyas, Matthew D Golub, David Sussillo, and Krishna V Shenoy. Computation through neural population dynamics. Annual review of neuroscience, 43(1):249–275, 2020. doi: 10.1146/annurev-neuro-092619-094115.

[16] Mark M. Churchland, John P. Cunningham, Matthew T. Kaufman, Justin D. Foster, Paul Nuyujukian, Stephen I. Ryu, Krishna V. Shenoy, and Krishna V. Shenoy. Neural population dynamics during reaching. Nature, 487(7405):51–56, 2012. doi: 10.1038/nature11129.

[17] Kristopher T Jensen, Naama Kadmon Harpaz, Ashesh K Dhawale, Steffen BE Wolff, and Bence P Ölveczky. Long-term stability of single neuron activity in the motor system. Nature neuroscience, 25(12):1664–1674, 2022. doi: 10.1038/s41593-022-01194-3.

[18] Juan A Gallego, Matthew G Perich, Raeed H Chowdhury, Sara A Solla, and Lee E Miller. Long-term stability of cortical population dynamics underlying consistent behavior. Nature neuroscience, 23(2):260–270, 2020. doi: 10.1038/s41593-019-0555-4.

[19] Javier F. Medina and Stephen G. Lisberger. Links from complex spikes to local plasticity and motor learning in the cerebellum of awake-behaving monkeys. Nature neuroscience, 11:1185 – 1192, 2008. URL https://api.semanticscholar.org/CorpusID:452344.

[20] Dylan J. Calame, Matthew I. Becker, and Abigail L. Person. Cerebellar associative learning underlies skilled reach adaptation. Nature Neuroscience, 26:1068–1079, 2021. URL https://api.semanticscholar.org/CorpusID:245367565.

[21] Amy J. Bastian. Understanding sensorimotor adaptation and learning for rehabilitation. Current Opinion in Neurology, 21(6):628, 2008. doi: 10.1097/WCO.0b013e328315a293. URL https://journals.lww.com/co-neurology/fulltext/2008/12000/understanding_sensorimotor_adaptation_and_learning.3.aspx.

[22] Jennifer L. Raymond and Javier F. Medina. Computational principles of supervised learning in the cerebellum. Annual Review of Neuroscience, 41:233–253, 2018. doi: 10.1146/annurev-neuro-080317-061948.

[23] N. Alex Cayco-Gajic and R. Angus Silver. Re-evaluating Circuit Mechanisms Underlying Pattern Separation. Neuron, 101(4):584–602, 2019. doi: 10.1016/j.neuron.2019.01.044. URL https://www.cell.com/neuron/abstract/S0896-6273(19)30071-6.

[24] Nuo Li and Thomas D Mrsic-Flogel. Cortico-cerebellar interactions during goal-directed behavior. Current opinion in neurobiology, 65:27–37, 2020.

[25] Zhenyu Gao, Courtney Davis, Alyse M. Thomas, Michael N. Economo, Amada M. Abrego, Karel Svoboda, Chris I. De Zeeuw, and Nuo Li. A cortico-cerebellar loop for motor planning. Nature, 563(7729):113–116, 2018. doi: 10.1038/s41586-018-0633-x.

[26] Mark J. Wagner, Tony Hyun Kim, Jonathan Kadmon, Nghia D. Nguyen, Surya Ganguli, Mark J. Schnitzer, and Liqun Luo. Shared cortex-cerebellum dynamics in the execution and learning of a motor task. Cell, 177(3):669–682.e24, 2019. doi: 10.1016/j.cell.2019.02.019.

[27] Joseph M. Galea, Alejandro Vazquez, Neel Pasricha, Jean-Jacques Orban de Xivry, and Pablo Celnik. Dissociating the Roles of the Cerebellum and Motor Cortex during Adaptive Learning: The Motor Cortex Retains What the Cerebellum Learns. Cerebral Cortex, 21(8):1761–1770, 2011. doi: 10.1093/cercor/bhq246. URL https://doi.org/10.1093/cercor/bhq246.

[28] Francois P Chabrol, Antonin Blot, and Thomas D Mrsic-Flogel. Cerebellar contribution to preparatory activity in motor neocortex. Neuron, 103(3):506–519, 2019.

[29] Rémi D. Proville, Maria Spolidoro, Nicolas Guyon, Guillaume P. Dugué, Fekrije Selimi, Philippe Isope, Daniela Popa, and Clément Léna. Cerebellum involvement in cortical sensorimotor circuits for the control of voluntary movements. Nature Neuroscience, 17(9):1233–1239, 2014. doi: 10.1038/nn.3773.

[30] Sharon Israely, Hugo Ninou, Ori Rajchert, Lee Elmaleh, Ran Harel, Firas Mawase, Jonathan Kadmon, and Yifat Prut. Cerebellar output shapes cortical preparatory activity during motor adaptation. Nat Commun, 16(1):2574, 2025. doi: 10.1038/s41467-025-57832-4. URL https://www.nature.com/articles/s41467-025-57832-4.

[31] Joseph Pemberton, Paul Chadderton, and Rui Ponte Costa. Cerebellar-driven cortical dynamics can enable task acquisition, switching and consolidation. Nat Commun, 15(1):10913, 2024. doi: 10.1038/s41467-024-55315-6. URL https://www.nature.com/articles/s41467-024-55315-6.

[32] James S Albus. A theory of cerebellar function. Mathematical Biosciences, 10(1-2):25–61, 1971. doi: 10.1016/0025-5564(71)90051-4.

[33] David C. Marr. A theory of cerebellar cortex. The Journal of Physiology, 202, 1969. URL https://api.semanticscholar.org/CorpusID:19906341.

[34] Masao Ito. Neurophysiological aspects of the cerebellar motor control system. International journal of neurology, 7 2:162–76, 1970. URL https://api.semanticscholar.org/CorpusID:19220808.

[35] M. Kawato, Kazunori Furukawa, and R. Suzuki. A hierarchical neural-network model for control and learning of voluntary movement. Biological Cybernetics, 57(3):169–185, 1987. doi: 10.1007/BF00364149.

[36] Mitsuo Kawato. Internal models for motor control and trajectory planning. Current Opinion in Neurobiology, 9:718–727, 1999. URL https://api.semanticscholar.org/CorpusID:878792.

[37] Anupama Sathyamurthy, Arnab Barik, Courtney I. Dobrott, Kaya J.E. Matson, Stefan Stoica, Randall Pursley, Alexander Theodore Chesler, and Ariel J. Levine. Cerebellospinal neurons regulate motor performance and motor learning. Cell reports, 31:107595 – 107595, 2020. URL https://api.semanticscholar.org/CorpusID:218634857.

[38] John F Kalaska, Stephen H Scott, Paul Cisek, and Lauren E Sergio. Cortical control of reaching movements. Current Opinion in Neurobiology, 7(6):849–859, 1997. doi: 10.1016/S0959-4388(97)80146-8. URL https://www.sciencedirect.com/science/article/pii/S0959438897801468.

[39] Joanna C. Chang, Matthew G. Perich, Lee E. Miller, Juan A. Gallego, and Claudia Clopath. De novo motor learning creates structure in neural activity that shapes adaptation. Nat Commun, 15(1):4084, 2024. doi: 10.1038/s41467-024-48008-7. URL https://www.nature.com/articles/s41467-024-48008-7.

[40] Harsha Gurnani and N Alex Cayco Gajic. Signatures of task learning in neural representations. Current Opinion in Neurobiology, 83:102759, 2023. ISSN 0959-4388. doi: 10.1016/j.conb.2023.102759. URL https://www.sciencedirect.com/science/article/pii/S0959438823000843.

[41] Hiroaki Gomi and Mitsuo Kawato. Neural network control for a closed-loop system using feedback-error-learning. Neural Networks, 6(7):933–946, 1993. doi: 10.1016/S0893-6080(09)80004-X.

[42] Aditya Gilra and Wulfram Gerstner. Predicting non-linear dynamics by stable local learning in a recurrent spiking neural network. eLife, 6:1–37, 2017. doi: 10.7554/eLife.28295.

[43] Barbara Feulner, M.G. Perich, Lee E. Miller, Claudia Clopath, and Juan A. Gallego. A neural implementation model of feedback-based motor learning. Nature Communications, 16, 2025. URL https://api.semanticscholar.org/CorpusID:276506680.

[44] Sophie Denéve, Alireza Alemi, and Ralph Bourdoukan. The brain as an efficient and robust adaptive learner. Neuron, 94(5):969–977, 2017. doi: 10.1016/j.neuron.2017.05.016.

[45] Daniel M. Wolpert, R. Chris Miall, and Mitsuo Kawato. Internal models in the cerebellum. Trends in Cognitive Sciences, 2:338–347, 1998. URL https://api.semanticscholar.org/CorpusID:10383815.

[46] Kurt Hornik, Maxwell B. Stinchcombe, and Halbert L. White. Multilayer feedforward networks are universal approximators. Neural Networks, 2:359–366, 1989. URL https://api.semanticscholar.org/CorpusID:2757547.

[47] Court Hull and Wade G. Regehr. The Cerebellar Cortex. Annual Review of Neuroscience, 45 (Volume 45, 2022):151–175, 2022. doi: 10.1146/annurev-neuro-091421-125115. URL https://www.annualreviews.org/content/journals/10.1146/annurev-neuro-091421-125115.

[48] Jia Zhu, Hana Hasanbegović, Liu D. Liu, Zhenyu Gao, and Nuo Li. Activity map of a corticocerebellar loop underlying motor planning. Nat Neurosci, 26(11):1916–1928, 2023. doi: 10.1038/s41593-023-01453-x. URL https://www.nature.com/articles/s41593-023-01453-x.

[49] Roberta M. Kelly and Peter L. Strick. Cerebellar loops with motor cortex and prefrontal cortex of a nonhuman primate. Journal of Neuroscience, 23(23):8432–8444, 2003. doi: 10.1523/jneurosci.23-23-08432.2003.

[50] Adriana Perez Rotondo, Dhruva V Raman, and Timothy O’Leary. How Cerebellar Architecture and Dense Activation Patterns Facilitate Online Learning in Dynamic Tasks. bioRxiv, page 2022.10.20.512268, 2023. doi: 10.1101/2022.10.20.512268.

[51] C. Asanuma, W. T. Thach, and E. G. Jones. Brainstem and spinal projections of the deep cerebellar nuclei in the monkey, with observations on the brainstem projections of the dorsal column nuclei. Brain Research Reviews, 5(3):299–322, 1983. doi: 10.1016/0165-0173(83)90017-6. URL https://www.sciencedirect.com/science/article/pii/0165017383900176.

[52] Clémentine Bosch-Bouju, Brian I. Hyland, and Louise C. Parr-Brownlie. Motor thalamus integration of cortical, cerebellar and basal ganglia information: Implications for normal and parkinsonian conditions. Front. Comput. Neurosci., 7, 2013. doi: 10.3389/fncom.2013.00163. URL https://www.frontiersin.org/journals/computational-neuroscience/articles/10.3389/fncom.2013.00163/full.

[53] Jake P. Stroud, Mason A. Porter, Guillaume Hennequin, and Tim P. Vogels. Motor primitives in space and time via targeted gain modulation in cortical networks. Nature neuroscience, 21:1774 – 1783, 2018. URL https://api.semanticscholar.org/CorpusID:53770571.

[54] David Sussillo, Mark M. Churchland, Matthew T. Kaufman, and Krishna V. Shenoy. A neural network that finds a naturalistic solution for the production of muscle activity. Nature Neuroscience, 18:1025–1033, 2015. URL https://api.semanticscholar.org/CorpusID:16351622.

[55] Ta-Chu Kao, Mahdieh Sadat Sadabadi, and Guillaume Hennequin. Optimal anticipatory control as a theory of motor preparation: A thalamo-cortical circuit model. Neuron, 109:1567 – 1581.e12, 2020. URL https://api.semanticscholar.org/CorpusID:214478881.

[56] Harsha Gurnani, Weixuan Liu, and BW Brunton. Feedback control of recurrent dynamics constrains learning timescales during motor adaptation. bioRxiv, 2024. URL https://api.semanticscholar.org/CorpusID:270068565.

[57] James B. Heald, Máté Lengyel and Daniel M. Wolpert. Contextual inference underlies the learning of sensorimotor repertoires. Nature, 600(7889):489–493, 2021. doi: 10.1038/s41586-021-04129-3. URL https://www.nature.com/articles/s41586-021-04129-3.

[58] Brandon J. Bhasin, Jennifer L. Raymond, and Mark S. Goldman. Synaptic weight dynamics underlying memory consolidation: Implications for learning rules, circuit organization, and circuit function. Proceedings of the National Academy of Sciences, 121(41):e2406010121, 2024. doi: 10.1073/pnas.2406010121. URL https://www.pnas.org/doi/10.1073/pnas.2406010121.

[59] Brendan A. Bicknell and Peter E. Latham. Fast and slow synaptic plasticity enables concurrent control and learning. eLife, 14, 2025. doi: 10.7554/eLife.105043.1. URL https://elifesciences.org/reviewed-preprints/105043.

[60] Risa Kawai, Timothy Markman, Rajesh Poddar, Raymond Ko, Antoniu L. Fantana, Ashesh K. Dhawale, Adam Raymond Kampff, and Bence P. Ölveczky. Motor cortex is required for learning but not for executing a motor skill. Neuron, 86:800–812, 2015. URL https://api.semanticscholar.org/CorpusID:14535811.

[61] Kristopher T. Jensen, Naama Kadmon Harpaz, Ashesh K. Dhawale, Steffen B. E. Wolff, and Bence P. Ölveczky. Long-term stability of single neuron activity in the motor system. bioRxiv, 2021. URL https://api.semanticscholar.org/CorpusID:240256118.

[62] Guang Yang, Feng Pan, and Wen-Biao Gan. Stably maintained dendritic spines are associated with lifelong memories. Nature, 462(7275):920–924, 2009. doi: 10.1038/nature08577. URL https://www.nature.com/articles/nature08577.

[63] Krishna V. Shenoy, Maneesh Sahani, and Mark M. Churchland. Cortical control of arm movements: a dynamical systems perspective. Annual review of neuroscience, 36:337–59, 2013. URL https://api.semanticscholar.org/CorpusID:17833963.

[64] Britton A. Sauerbrei, Jian-Zhong Guo, Jeremy D. Cohen, Matteo Mischiati, Wendy Guo, Mayank Kabra, Nakul Verma, Brett D. Mensh, Kristin Branson, and Adam W. Hantman. Cortical pattern generation during dexterous movement is input-driven. Nature, 577:386 – 391, 2019. URL https://api.semanticscholar.org/CorpusID:209481603.

[65] Daniel J. O’Shea, Lea Duncker, Werapong Goo, Xulu Sun, Saurabh Vyas, Eric M. Trautmann, Ilka Diester, Charu Ramakrishnan, Karl Deisseroth, Maneesh Sahani, and Krishna V. Shenoy. Direct neural perturbations reveal a dynamical mechanism for robust computation, 2022. URL https://www.biorxiv.org/content/10.1101/2022.12.16.520768v1.

[66] Emily R. Oby, Alan D. Degenhart, Erinn M. Grigsby, Asma Motiwala, Nicole T. McClain, Patrick J. Marino, Byron M. Yu, and Aaron P. Batista. Dynamical constraints on neural population activity. Nat Neurosci, 28(2):383–393, 2025. doi: 10.1038/s41593-024-01845-7. URL https://www.nature.com/articles/s41593-024-01845-7.

[67] Ya-weng Tseng, Jörn Diedrichsen, John W. Krakauer, Reza Shadmehr, and Amy J. Bastian. Sensory Prediction Errors Drive Cerebellum-Dependent Adaptation of Reaching. Journal of Neurophysiology, 98(1):54–62, 2007. doi: 10.1152/jn.00266.2007.

[68] D. Timmann, J. Drepper, M. Frings, M. Maschke, S. Richter, M. Gerwig, and F. P. Kolb. The human cerebellum contributes to motor, emotional and cognitive associative learning. A review. Cortex, 46(7):845–857, 2010. doi: 10.1016/j.cortex.2009.06.009. URL https://www.sciencedirect.com/science/article/pii/S0010945209002044.

[69] Reza Shadmehr, Maurice A. Smith, and John W. Krakauer. Error Correction, Sensory Prediction, and Adaptation in Motor Control. Annual Review of Neuroscience, 33(Volume 33, 2010):89–108, 2010. doi: 10.1146/annurev-neuro-060909-153135.

[70] David J. Herzfeld, Yoshiko Kojima, Robijanto Soetedjo, and Reza Shadmehr. Encoding of error and learning to correct that error by the Purkinje cells of the cerebellum. Nat Neurosci, 21(5):736–743, 2018. doi: 10.1038/s41593-018-0136-y. URL https://www.nature.com/articles/s41593-018-0136-y.

[71] Shogo Ohmae and Javier F. Medina. Climbing fibers encode a temporal-difference prediction error during cerebellar learning in mice. Nat Neurosci, 18(12):1798–1803, 2015. doi: 10.1038/nn.4167. URL https://www.nature.com/articles/nn.4167.

[72] Brenda D. Houck and Abigail L. Person. Cerebellar loops: A review of the nucleocortical pathway. The Cerebellum, 13:378 – 385, 2013. URL https://api.semanticscholar.org/CorpusID:913782.

[73] Narender Ramnani. The primate cortico-cerebellar system: Anatomy and function. Nat Rev Neurosci, 7(7):511–522, 2006. doi: 10.1038/nrn1953. URL https://www.nature.com/articles/nrn1953.

[74] J. A. Menéndez, J. A. Hennig, M. D. Golub, E. R. Oby, P. T. Sadtler, A. P. Batista, S. M. Chase, B. M. Yu, and P. E. Latham. A theory of brain-computer interface learning via low-dimensional control, 2024. URL http://biorxiv.org/lookup/doi/10.1101/2024.04.18.589952.

[75] Samuel D. McDougle, Richard B. Ivry, and Jordan A. Taylor. Taking Aim at the Cognitive Side of Learning in Sensorimotor Adaptation Tasks. Trends in Cognitive Sciences, 20(7):535– 544, 2016. doi: 10.1016/j.tics.2016.05.002. URL https://www.sciencedirect.com/science/article/pii/S1364661316300419.

[76] Ellen Boven, Joseph Pemberton, Paul Chadderton, Richard Apps, and Rui Ponte Costa. Cerebro-cerebellar networks facilitate learning through feedback decoupling. Nature Communications, 14(1):1–18, 2023. doi: 10.1038/s41467-022-35658-8. URL https://www.nature.com/articles/s41467-022-35658-8.

[77] Laureline Logiaco, L. F. Abbott, and Sean Escola. Thalamic control of cortical dynamics in a model of flexible motor sequencing. Cell reports, 35:109090 – 109090, 2021. URL https://api.semanticscholar.org/CorpusID:235320723.

[78] Diego E. Aldarondo, Josh Merel, Jesse D. Marshall, Leonard Hasenclever, Ugne Klibaite, Amanda Gellis, Yuval Tassa, Greg Wayne, Matthew M. Botvinick, and Bence P. Ölveczky. A virtual rodent predicts the structure of neural activity across behaviors. Nature, 2024. URL https://api.semanticscholar.org/CorpusID:270389836.

[79] Guy Bouvier, Johnatan Aljadeff, Claudia Clopath, Célian Bimbard, Jonas Ranft, Antonin Blot, Jean-Pierre Nadal, Nicolas J.-B. Brunel, Vincent Hakim, and Boris Barbour. Cerebellar learning using perturbations. eLife, 7, 2016. URL https://api.semanticscholar.org/CorpusID:10765994.

[80] Dimitar Kostadinov and Michael Häusser. Reward signals in the cerebellum: Origins, targets, and functional implications. Neuron, 110(8):1290–1303, 2022. doi: 10.1016/j.neuron.2022.02.015. URL https://www.cell.com/neuron/abstract/S0896-6273(22)00180-5.

[81] Court Hull. Prediction signals in the cerebellum: Beyond supervised motor learning. eLife, 9:e54073, 2020. doi: 10.7554/eLife.54073.

[82] Elisa Galliano, Zhenyu Gao, Martijn Schonewille, Boyan Todorov, Esther Simons, Andreea S. Pop, Egidio D’Angelo, Arn M.J.M. van den Maagdenberg, Freek E. Hoebeek, and Chris I. De Zeeuw. Silencing the Majority of Cerebellar Granule Cells Uncovers Their Essential Role in Motor Learning and Consolidation. Cell Reports, 3(4):1239–1251, 2013. doi: 10.1016/j.celrep.2013.03.023. URL https://www.sciencedirect.com/science/article/pii/S2211124713001307.

[83] Nicolas Catz, Peter W. Dicke, and Peter Thier. Cerebellar Complex Spike Firing Is Suitable to Induce as Well as to Stabilize Motor Learning. Current Biology, 15(24):2179–2189, 2005. doi: 10.1016/j.cub.2005.11.037. URL https://www.sciencedirect.com/science/article/pii/S0960982205014466.

[84] Henry H. Yin, Shweta Prasad Mulcare, Monica R. F. Hilário, Emily Clouse, Terrell Holloway, Margaret I. Davis, Anita C. Hansson, David M. Lovinger, and Rui M. Costa. Dynamic reorganization of striatal circuits during the acquisition and consolidation of a skill. Nat Neurosci, 12(3): 333–341, 2009. doi: 10.1038/nn.2261. URL https://www.nature.com/articles/nn.2261.

[85] Robert M. French. Catastrophic forgetting in connectionist networks. Trends in Cognitive Sciences, 3:128–135, 1999. URL https://api.semanticscholar.org/CorpusID:2691726.

[86] Lea Duncker, Laura Driscoll, Krishna V Shenoy, Maneesh Sahani, and David Sussillo. Organizing recurrent network dynamics by task-computation to enable continual learning. In Advances in Neural Information Processing Systems, volume 33, pages 14387–14397. Curran Associates, Inc., 2020. URL https://proceedings.neurips.cc/paper/2020/hash/a576eafbce762079f7d1f77fca1c5cc2-Abstract.html.

[87] Xulu Sun, Daniel J. O’Shea, Matthew D. Golub, Eric M. Trautmann, Saurabh Vyas, Stephen I. Ryu, and Krishna V. Shenoy. Cortical preparatory activity indexes learned motor memories. Nature, 602(7896):274–279, 2022. doi: 10.1038/s41586-021-04329-x. URL https://www.nature.com/articles/s41586-021-04329-x.

[88] Adam Paszke, Sam Gross, Francisco Massa, Adam Lerer, James Bradbury, Gregory Chanan, Trevor Killeen, Zeming Lin, Natalia Gimelshein, Luca Antiga, Alban Desmaison, Andreas Köpf, Edward Yang, Zach DeVito, Martin Raison, Alykhan Tejani, Sasank Chilamkurthy, Benoit Steiner, Lu Fang, Junjie Bai, and Soumith Chintala. Pytorch: An imperative style, high-performance deep learning library. ArXiv, abs/1912.01703, 2019. URL https://api.semanticscholar.org/CorpusID:202786778.

[89] Kaiming He, Xiangyu Zhang, Shaoqing Ren, and Jian Sun. Delving deep into rectifiers: Surpassing human-level performance on imagenet classification, 2015. URL https://arxiv.org/abs/1502.01852.

[90] Xavier Glorot and Yoshua Bengio. Understanding the difficulty of training deep feedforward neural networks. In International Conference on Artificial Intelligence and Statistics, 2010. URL https://api.semanticscholar.org/CorpusID:5575601.

[91] Diederik P. Kingma and Jimmy Ba. Adam: A method for stochastic optimization. CoRR, abs/1412.6980, 2014. URL https://api.semanticscholar.org/CorpusID:6628106.

