## Supplementary Material for "Distributed learning across fast and slow neural systems supports efficient motor adaptation"

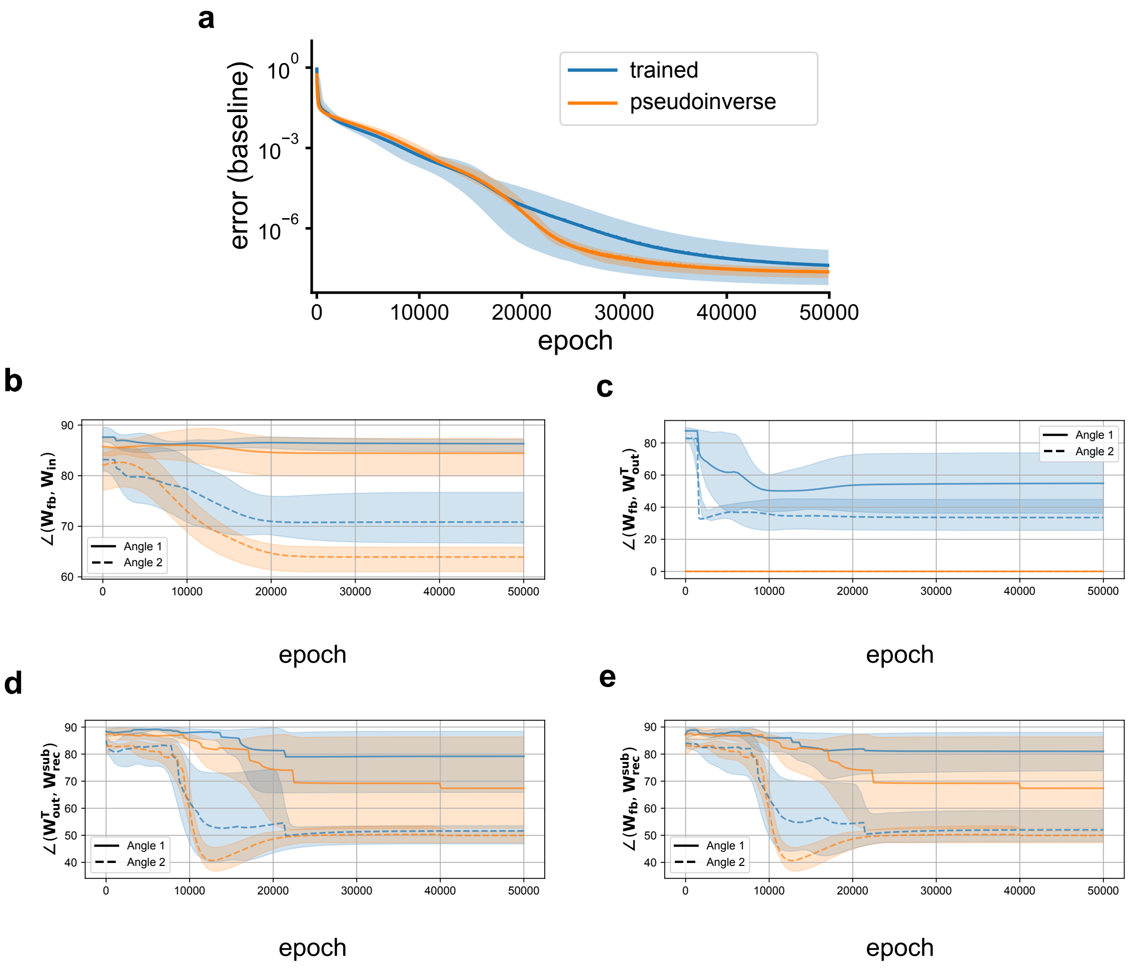


**Supplementary Figure 1: Pre-training metrics.** **a**, Performance of feedback controller RNNs during pretraining of the baseline center-out reach task. We compare models with output and feedback weights trained via backpropagation (blue) with models in which $W_{fb}$ and $W_{out}$ were fixed (not trained), setting $W_{fb} = W_{out}^{+}$ (orange). Although fewer parameters were being optimized, fixing output and feedback weights improved convergence. $W_{rec}$ and $W_{in}$ were trained in both cases. **b-d**, Evolution of alignment between principal eigenvectors of different weight matrices. **b**, Alignment between input and feedback weights (principal angles between the column spaces of $W_{in}$ and $W_{fb}$). **c**, Alignment between the feedback and output weights (principal angles between the column space of $W_{fb}$ and row space of $W_{out}$). **d**, Alignment between the recurrent weights and output weights (angles of the subspace corresponding to the top two eigenvalues of $W_{rec}$ and the rows of $W_{out}$). **e**, Alignment between the recurrent and feedback weights (angles of the subspace corresponding to the top two eigenvalues of $W_{rec}$ and the columns of $W_{fb}$).

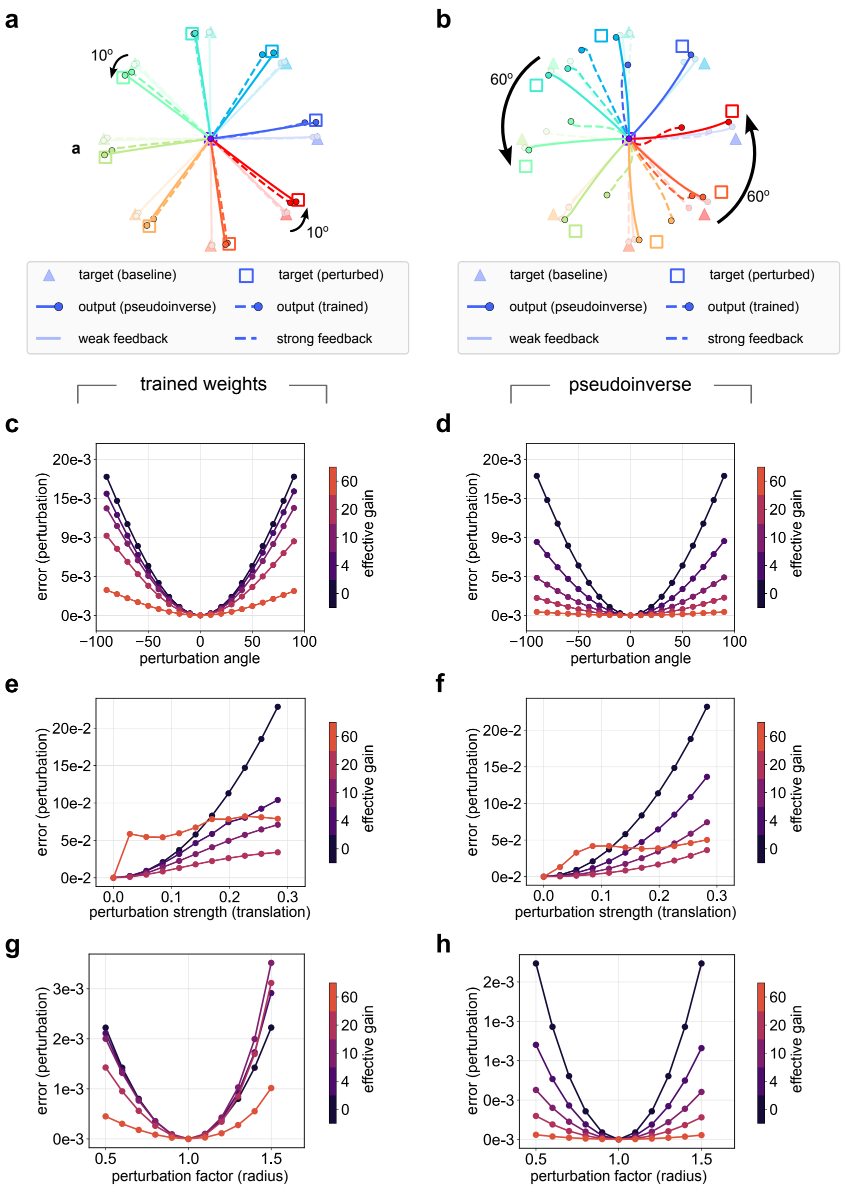


**Supplementary Figure 2: Performance of pre-trained feedback controllers under perturbation (fixed weights). a-b**, Hand trajectories under 10° (**a**) and 60° (**b**) rotational perturbations. Strong/weak feedback trajectories (dark/light trajectories) correspond to feedback gains of $\kappa = 30$ and $\kappa = 1$, respectively. Dashed/solid lines indicate trained/pseudoinverse feedback weights, respectively. **c-h**, Systematic tests of feedback controllers with backprop-trained (left column) and fixed pseudoinverse weights (right column) under different perturbations types and magnitudes, for different values of feedback strength ("effective gain"). **c-d**, Rotations of target trajectories. Rotation angle is reported in degrees. **e-f**, Translations of target endpoints. We tested 11 evenly spaced translations along the diagonal from [0,0] to [–0.2,–0.2]. The perturbation strength is the Euclidean distance from the unperturbed center of the trajectories. **g-h**, Radius changes of target endpoints. Starting from the original circular set of endpoints ($\mathbf{Y}_b$, see Methods), new trajectories were generated by scaling the radius of the circle by the reported perturbation factors.

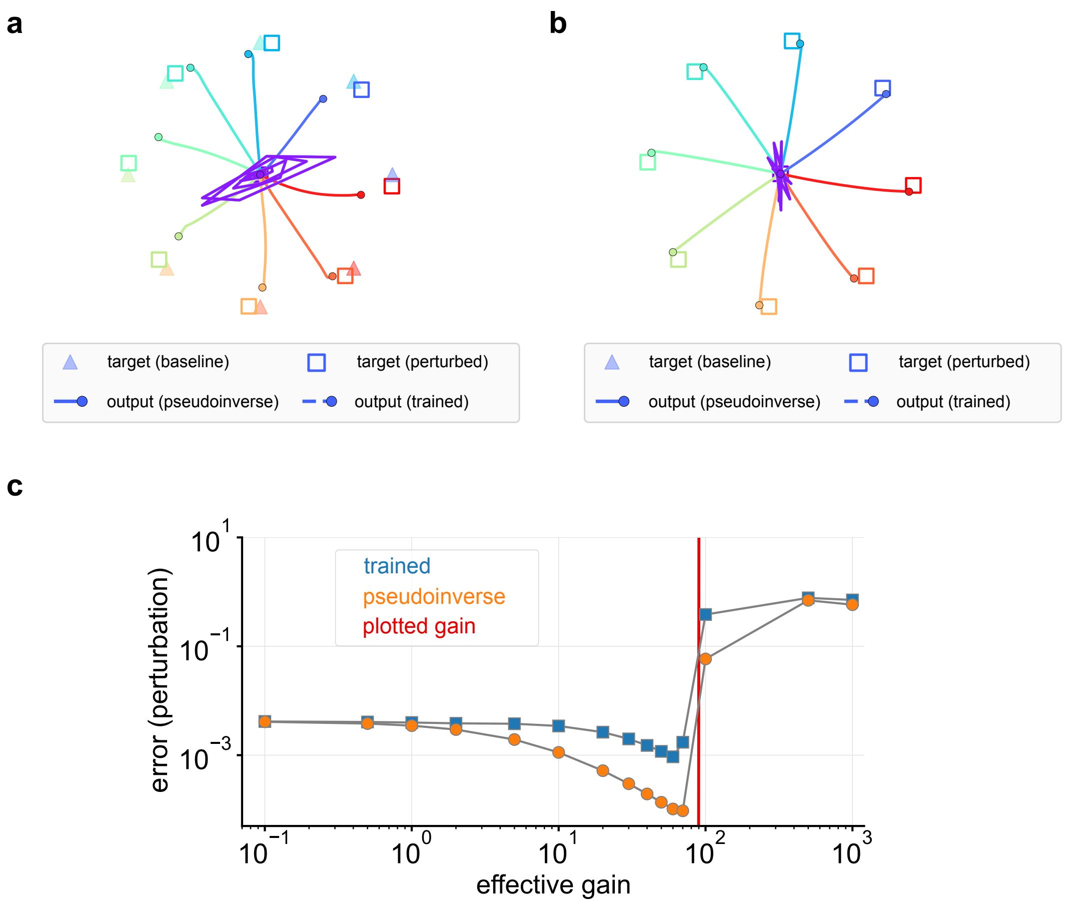


**Supplementary Figure 3: Perturbation error and unstable trajectories. a-b**, Output trajectories in response to a 40° rotational perturbations for feedback controllers with backprop-trained (**a**) and fixed pseudoinverse feedback weights (**b**), when fixing the effective gain to $\simeq 90$ (red line in **c**; equivalent to $\kappa = 45$). **c**, Error during the perturbation, showing a loss of stability at effective gains above $\sim 60$.

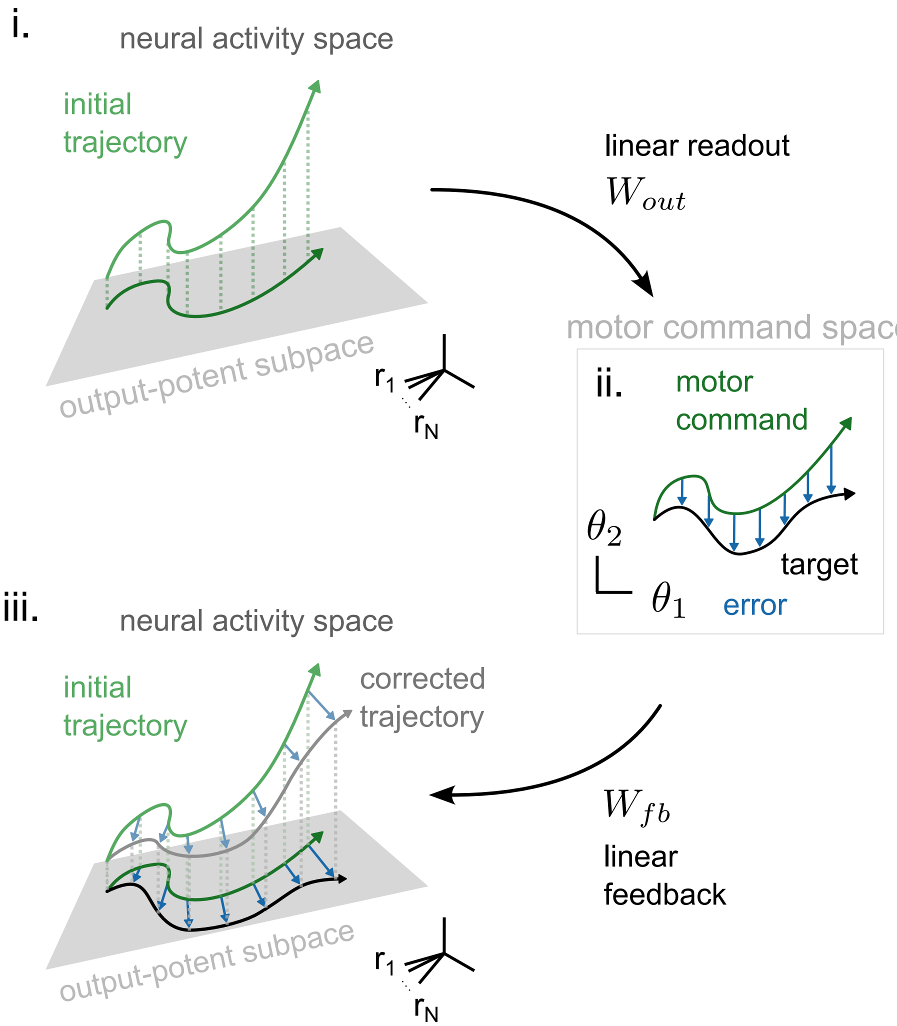


**Supplementary Figure 4: The pseudoinverse feedback mapping**, Schematic illustrating the geometric intuition behind the pseudoinverse feedback weights. *i.* The initial (uncorrected) controller trajectory (green) in an $n$-dimensional neural space is mapped to the 2-dimensional motor command space via the readout weights $W_{out}$. Because the neural space is higher-dimensional than the motor command space, only changes aligned with the row space of $W_{out}$ (the output-potent subspace, grey) will affect behavior. *ii.* The 2-dimensional error signal (blue) is the residual between the target (black) and the actual (green) motor command. *iii.* The controller trajectory is corrected by projecting the error back through the pseudoinverse $W_{fb} = W_{out}^{+}$, ensuring correct alignment of this feedback within the output-potent subspace.

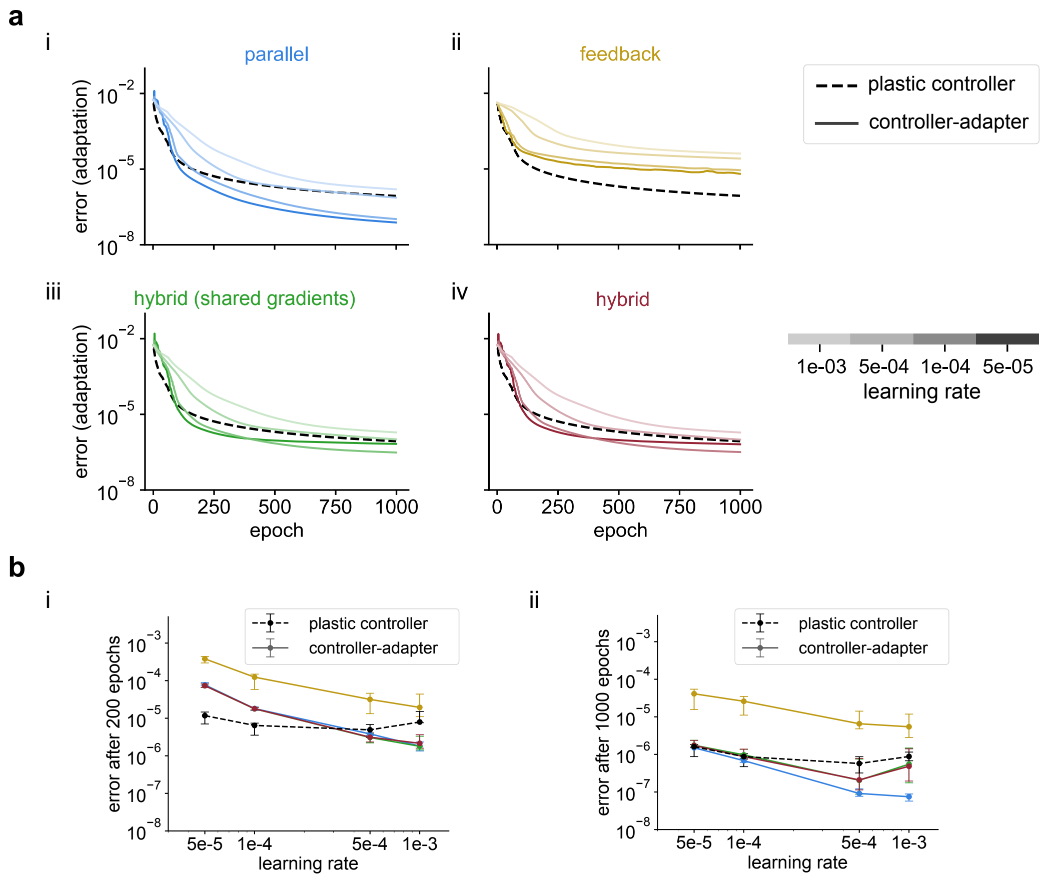


**Supplementary Figure 5: Performance of different controller-adapter architectures.** **a**, Here, we show the error during the adaptation task for each specific architecture described for the controller-adapter networks (parallel, feedback, and hybrid, with controller weights fixed), for different learning rates. Different shades correspond to different learning rates, according to the legend. *iii-iv* contain results for the hybrid model with (*iii*) and without (*iv*) shared gradients between the controller and the adapter (see Methods). **b** Average MSE after 200 (*i*) and 1000 epochs (*i*) for each architecture and learning rates in **a** (same color-coding). Error bars correspond to min-max values over 5 seeds, and each point of the line is the median value.

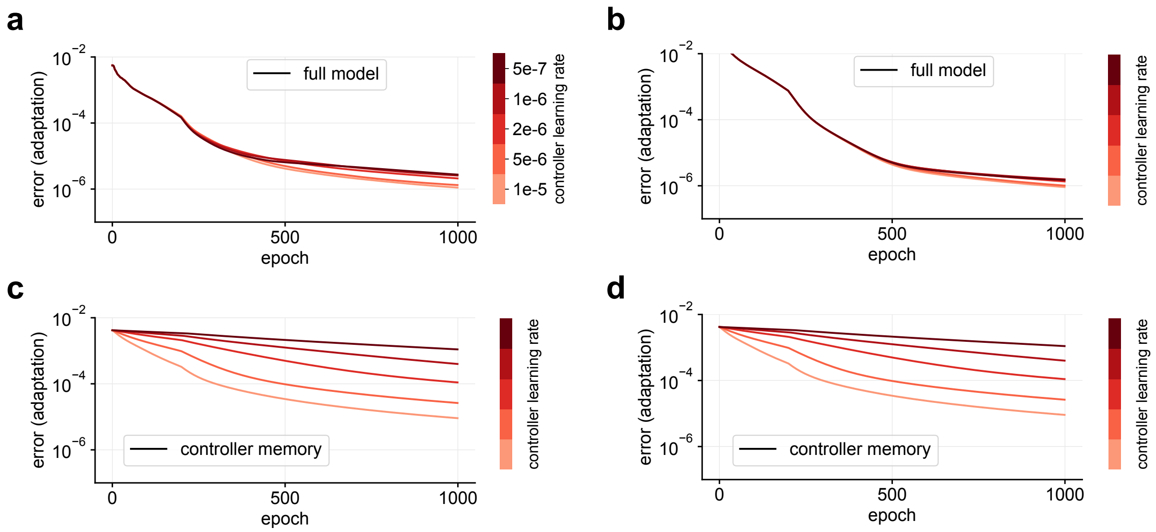


**Supplementary Figure 6: Performance of the full model and the controller memory under** $40°$ **rotation adaptation (backprop-trained controller).** Here, we show the error of the hybrid architecture during the standard adaptation task, for different values of the controller learning rate, while varying the controller learning rate strength (shade of red). **a-b**, Evolution of the full model error for weak (**a**, $\kappa = 1$) and strong (**b**, $\kappa = 20$) feedback strength regimes. **c-d**, Consolidation of the controller memory for weak and strong feedback, respectively. The adapter learning rate was set to 1e-4. Same color legend for all panels.

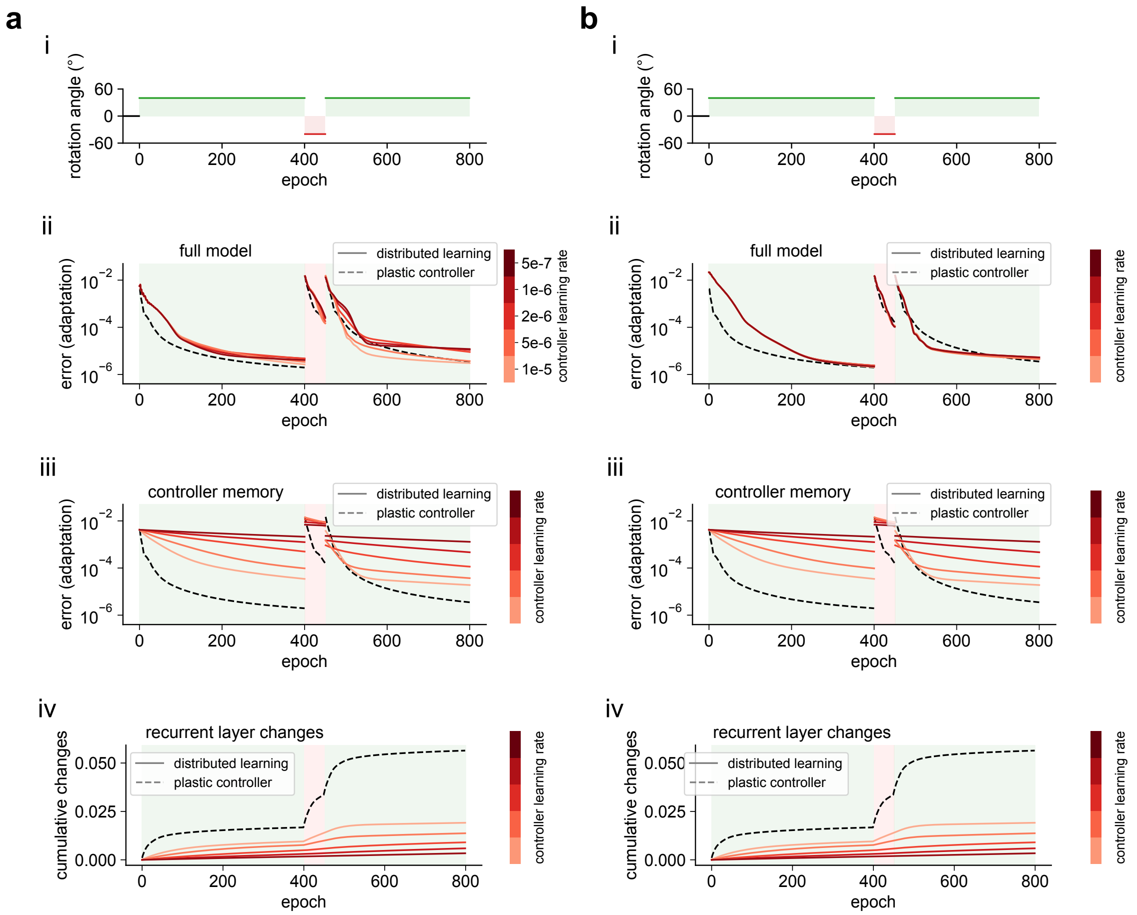


**Supplementary Figure 7: Performance of the distributed learning model on a re-adaptation task (backprop-trained controller).** **a**, Weak feedback regime ($\kappa = 1$). *i*, Schematic of the task. *ii*, The model shows improved savings (faster relearning with respect to the initial learning phase, see Main text) for faster controller memory consolidation speeds (larger learning rate). The single module plastic controller (dashed black line) showed no savings. *iii*, Savings can be explained by the quick error drop of the decoded controller memory once the initial rotation is restored (onset of second green region). *iv*, Cumulative changes in the recurrent weights. The plastic controller showed consistently larger changes in the recurrent weights compared to the distributed learning models. **b**, Same as **a** for the strong feedback regime ($\kappa = 20$). We note that in this case, the controller learning rate has no impact on full model performance (cf. **b**, *ii* with **a**, *ii*). Adapter learning rate is fixed to 2e-4. Same color legend for all panels.

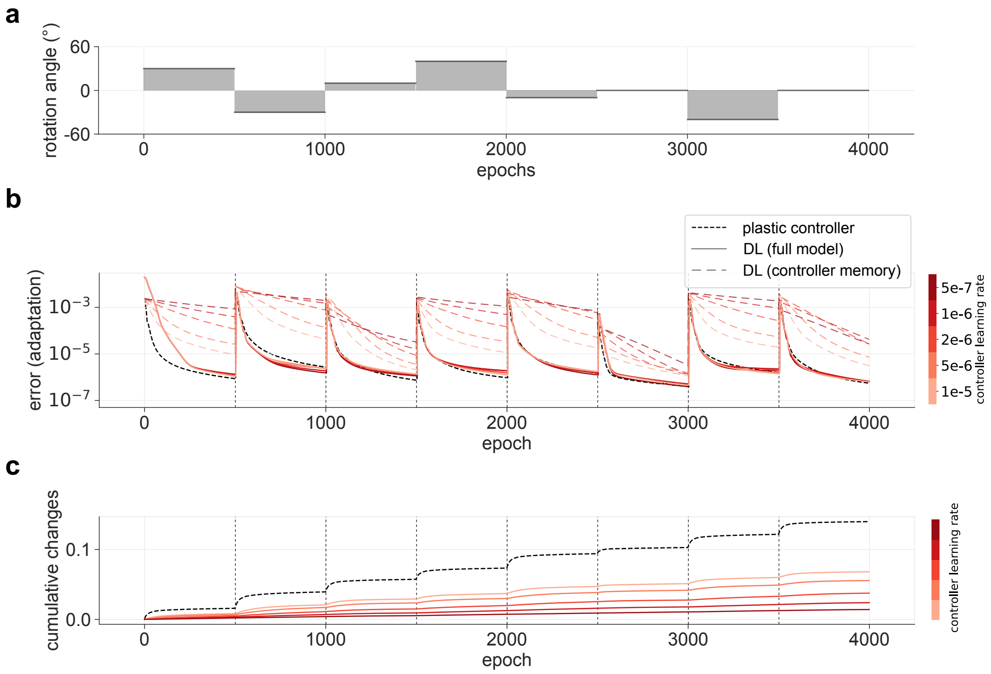


**Supplementary Figure 8: Distributed learning allows memory preservation on a continual learning task (backprop-trained controller).** Performance of the distributed learning model (DL) to repeated perturbations (500 epochs each). **a**, A continual learning task consisting of a sequence of random perturbations varying rotation angle, followed by a final washout phase (no rotation). **b**, Performance of the model with different controller learning rates (shade of red). A plastic controller is shown for reference (dashed black). All models showed comparable performance (solid lines) but vastly different memory consolidation speeds (colored dashed lines, showing consolidation of the controller memory). **c**, Cumulative changes of the controller's recurrent weights throughout the continual learning task. Compared to the plastic controller (black), all distributed learning models (red, color-coded according to learning rate) show strong reduction of changes of the recurrent dynamics. Adapter learning rate is fixed to 2e-4. Same color legend in **b** and **c**.

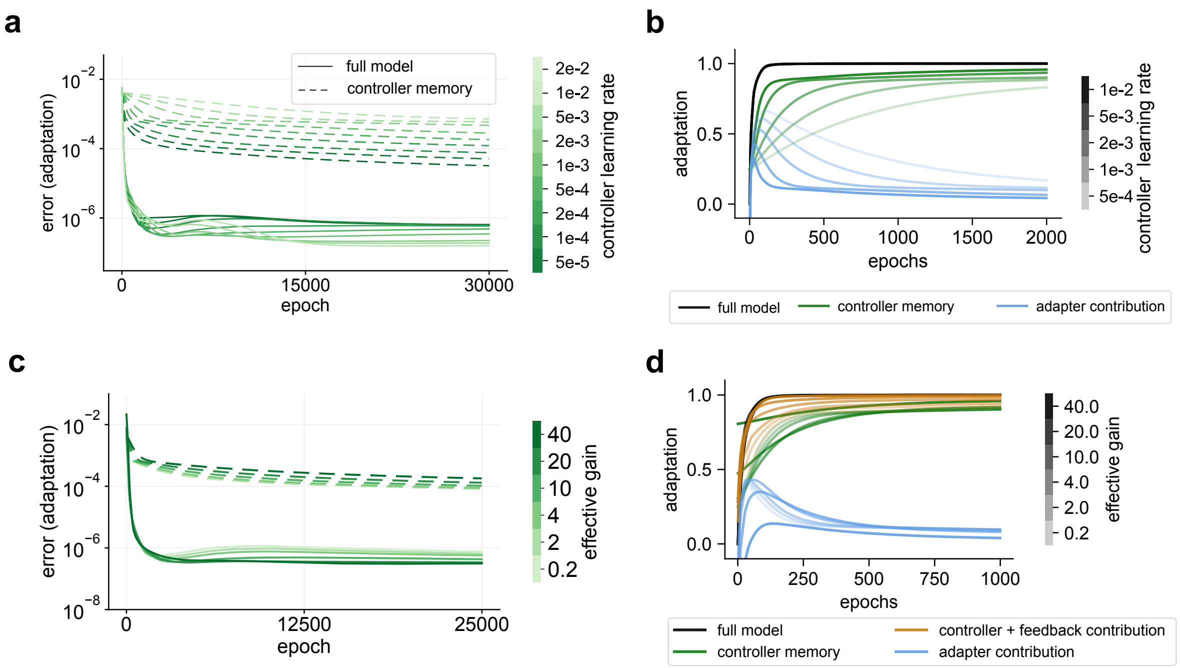


**Supplementary Figure 9: Varying contributions of the adapter and controller (local rule).** **a**, Performance of the distributed learning model over learning, for different values of the controller learning rate (shades of green). Solid lines correspond to full model error, while dashed lines represent controller memory error. **b**, Contributions of the controller and adapter to adaptation. Shade corresponds to different local learning rate in the controller. **c-d**, Effect of feedback strength (as effective gain) over learning. Stronger feedback led to better convergence of full model error, but results in slower consolidation. Controller learning rate is 5e-3. **c**, Performance of the full model (solid) and decoded controller memory (dashed), color coded by effective gain with different shades of green. **d**, Contributions of the adapter (blue), controller memory (green), and controller + feedback (gold). In all panels, the adapter learning rate is fixed to 1e-4.

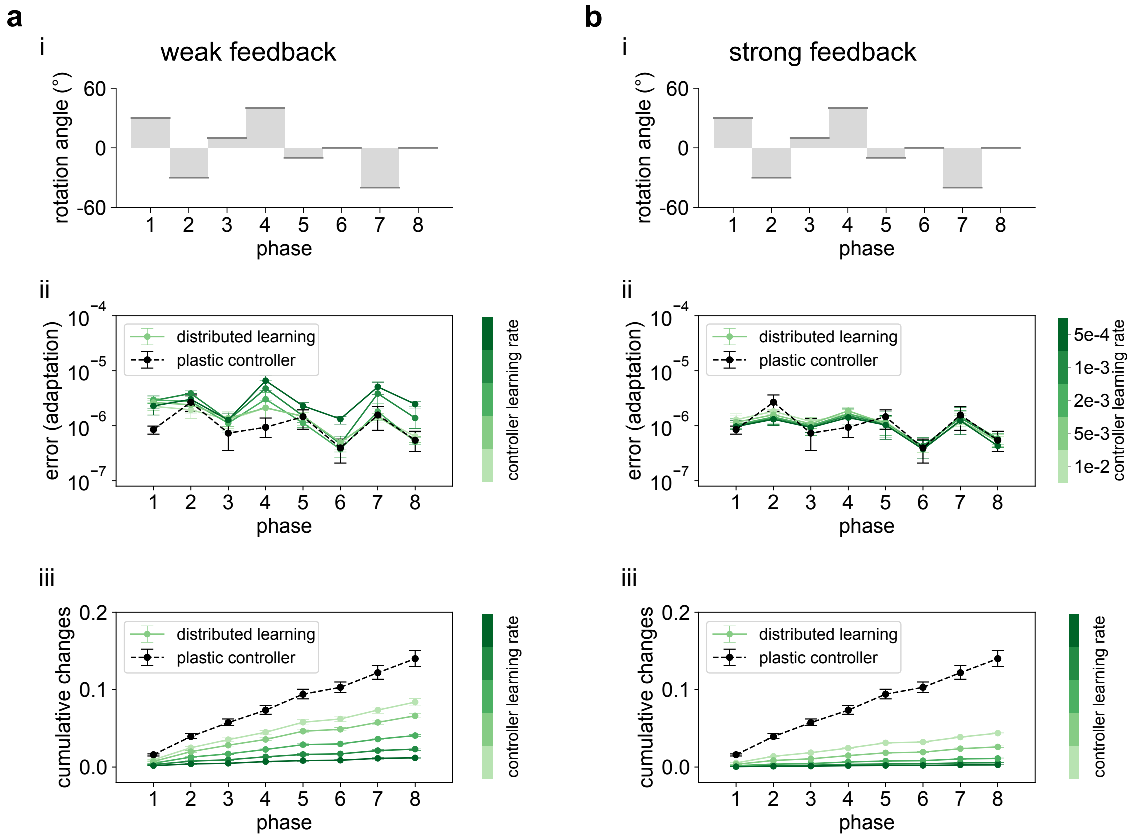


**Supplementary Figure 10: Performance of distributed learning model on a continual learning task (local rule).** **a,** Weak feedback regime ($\kappa = 1$). *i*, Schematic of the task. *ii*, Adaptation error at the end of each phase. *iii*, Cumulative changes over the recurrent layer. Solid lines shows distributed learning model performance, for different controller learning rates (shades of green). A plastic controller is shown in black. Error bars shows median (dots) and min-max values over 5 different seeds. Adapter learning rate is fixed at 2e-4. **b**, Same as **a** for the strong feedback regime ($\kappa = 20$). Strong feedback largely improves convergence of the distributed learning model across controller learning rates (cf. green lines in **a**, *ii* and **b**,*ii*). Same color legend for all panels.

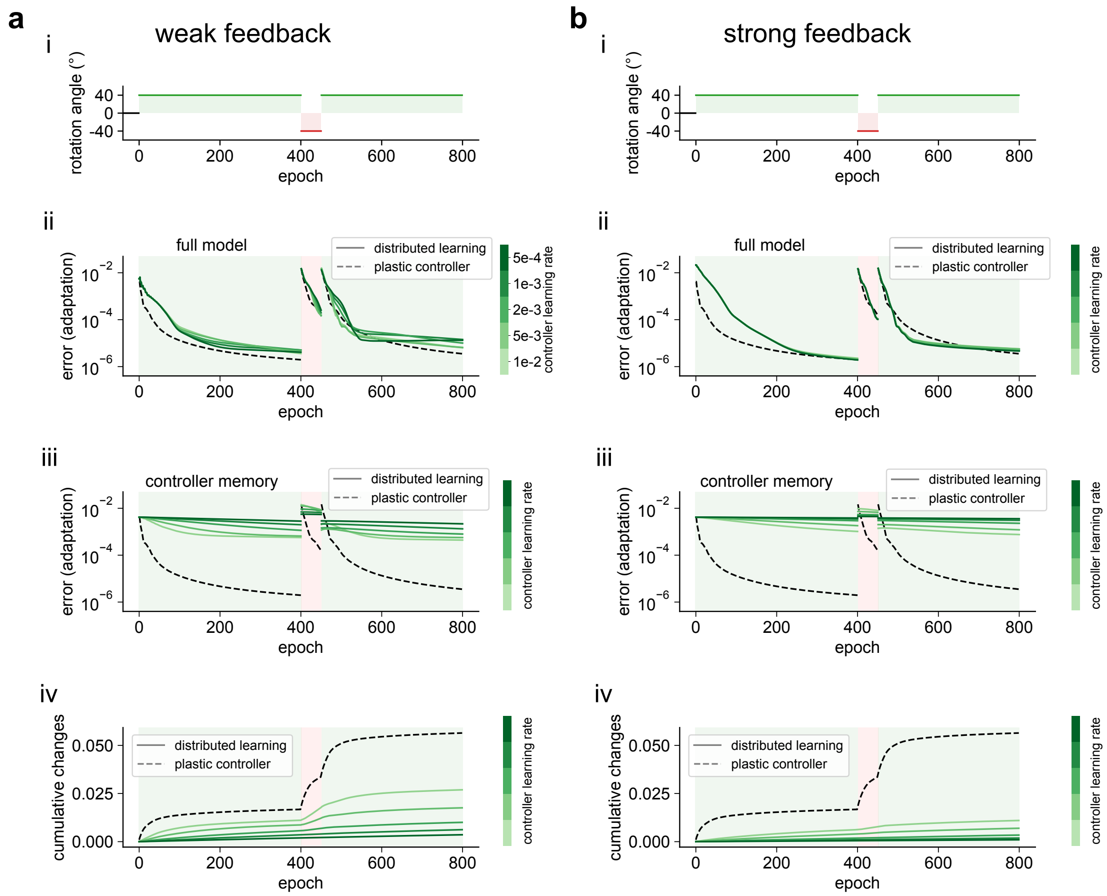


**Supplementary Figure 11: Performance of the distributed learning model on a re-adaptation task (local rule).** **a**, Weak feedback regime ($\kappa = 1$). *i*, Schematic of the task. *ii*, As for the backprop-consolidation case (Supplementary Figure 7), savings can be observed in the rapid error drop in the second 40° adaptation phase. The single module plastic controller (dashed black line) showed no savings. *iii*, Consolidation of the controller memory. *iv*, A plastic controller showed consistently larger cumulative changes over the recurrent weights compared to the distributed learning models. **b**, Same as **a** for the strong feedback regime. We note that, in this case, controller consolidation speed has no impact on full model performance, re-adaptation becomes faster than the plastic controller case (cf. **a**, *ii* with **b**, *ii*), and cumulative changes in the controller recurrent weights decreased (*iii*, cf. **a**, *iv* with **b**, *iv*). Adapter learning rate is fixed to 2e-4. Same color legend for all panels.

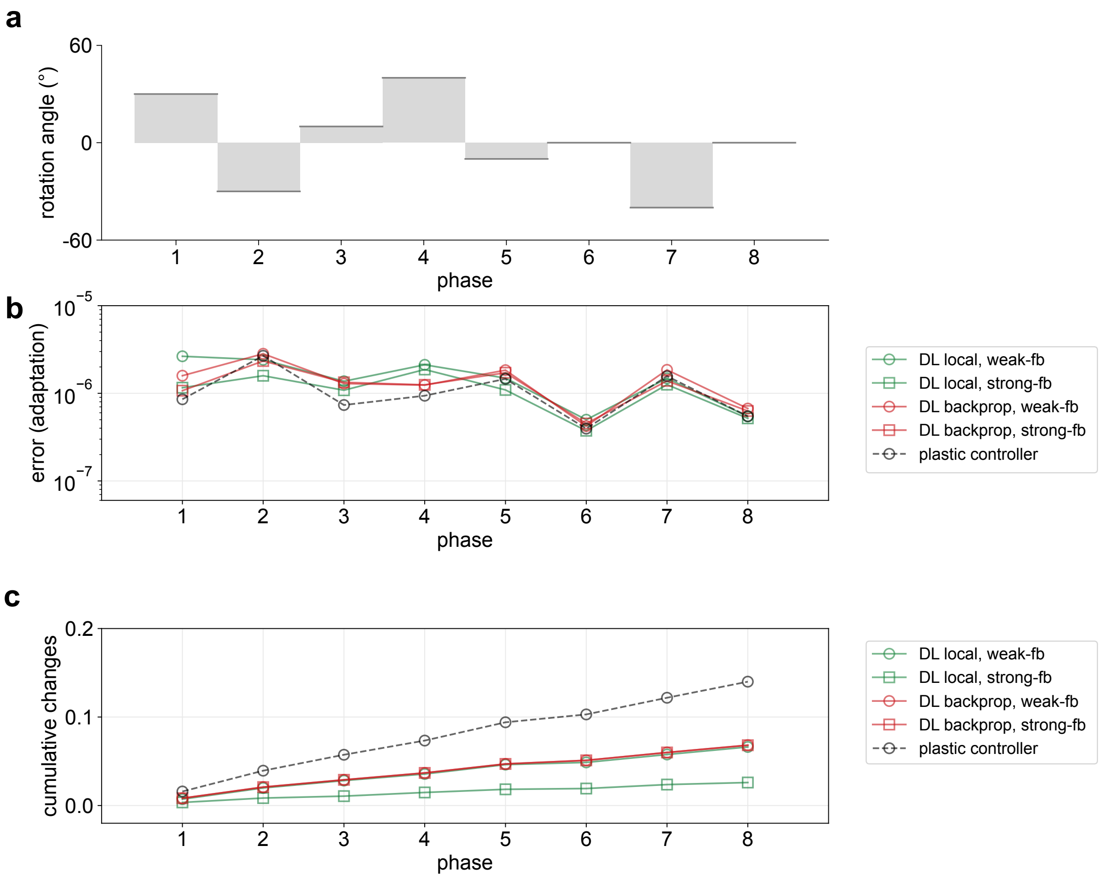


**Supplementary Figure 12: Comparison of different models in the continual learning task.** **a**, Schematic of the task. **b**, Adaptation error of each model at the end of each phase (median over 5 different seeds). Solid lines correspond to the distributed learning models (DL), with backprop (red) or local rule (green) consolidation. For each case, both weak (circles) and strong (squares) feedback regimes are shown. Dashed black line corresponds to the plastic controller. **c**, Cumulative changes of the controller's recurrent weights.

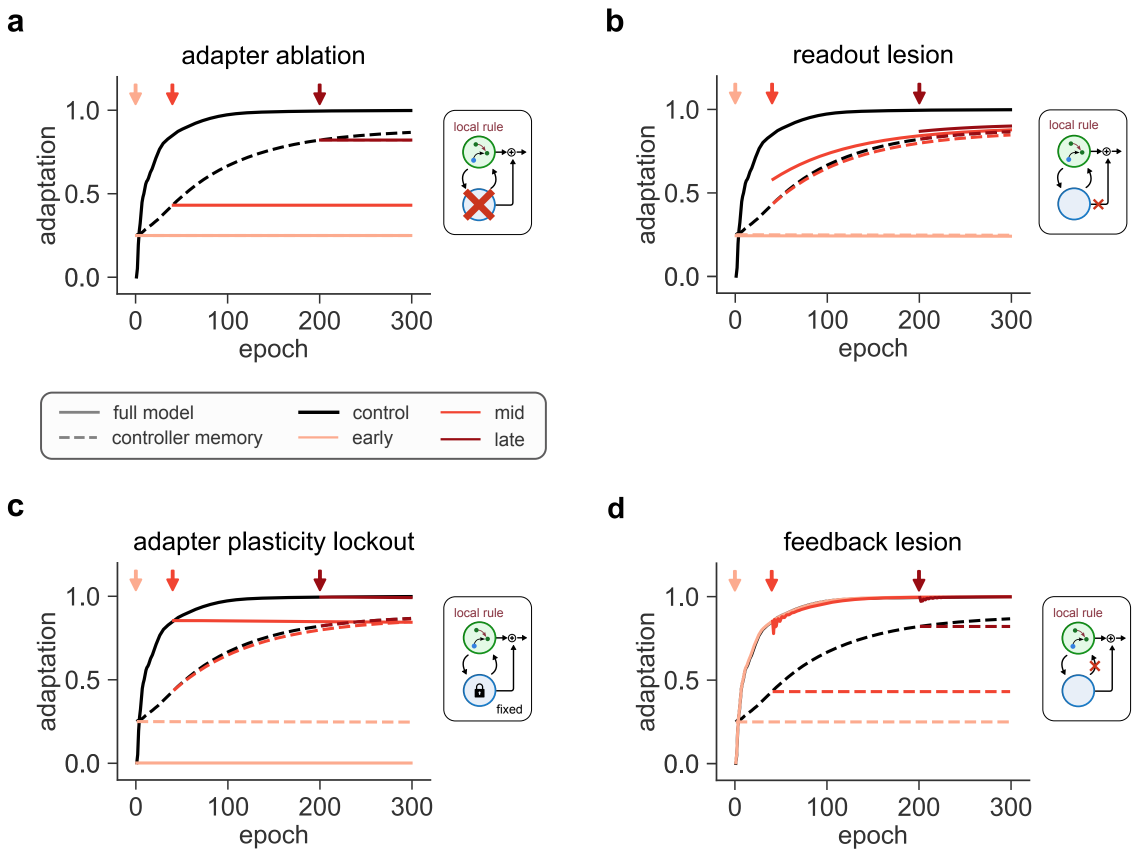


**Supplementary Figure 13: Signatures of distributed learning with tutored consolidation (local rule).** Adaptation metrics for different ablation tests of the full distributed learning model. For each panel, a specific component of the model was inactivated during learning, at either early (light red), mid (red) or late (dark red) stages of learning. For reference, the control (no ablation) is shown in black. Solid lines indicate the performance of the full model. Dashed black lines indicate the controller memory consolidation in the control (no ablation), while dashed red lines indicate the controller memory consolidation after ablation. All simulations used controller local learning rate 5e-3. **a**, When fully ablating the adapter network, performance drops to the level that the controller had consolidated at damage onset. No further learning occurs, due to the absence of teaching signal for the remaining module, the controller. **b**, Selectively lesioning the adapter readout projection caused an initial drop in performance. However, the full network still continued to learn, thanks to the preservation of the adapter feedback teaching signal (mid and late inactivations). Note that the performance at lesion onset (especially for the mid-learning inactivation) is higher than the controller memory due to the contribution of the adapter feedback to the controller, which is left intact (in contrast to **a**). **c**, The importance of the teaching signal could be observed by simply freezing the adapter weights, which led to a plateau in learning without an initial drop in performance as the adapter pathways were left intact. Interestingly, in mid and late interventions, controller consolidation continued to improve even if there was no change in the performance of the full network (dashed red lines), as the adapter had learned a sufficiently accurate error prediction signal that was able to guide controller plasticity. **d**, Conversely, lesioning the adapter feedback had little effect on full network performance as the adapter readout was able to fully fine-tune the controller output. However, the controller was no longer able to consolidate its memories.
